# Computational Design of Two Novel BRAF V600E Inhibitors: Exploiting Sulfoximine Bioisosterism and Chiral Constraints to Evade Paradoxical Activation

**DOI:** 10.64898/2026.06.25.734343

**Authors:** Zoe H. Yu, Emma R. Morrow, Justin B. Siegel

**Affiliations:** Department of Chemistry, University of California, Davis, Davis, California, United States of America; Genome Center, University of California, Davis, Davis, California, United States of America; Department of Biochemistry and Molecular Medicine, University of California, Davis, Davis, California, United States of America

## Abstract

Metastatic melanoma is an aggressive cutaneous malignancy frequently driven by the oncogenic V600E mutation within the BRAF kinase. While first-generation Type IS BRAF inhibitors, such as dabrafenib, are currently prescribed to target this specific molecular vulnerability, paradoxical MAPK pathway activation, and acquired drug resistance necessitate the continuous development of structurally optimized lead molecules. In this study, chemical intuition, bioisosteric replacement, and computational molecular docking were employed to propose two novel *BRAF*^*V600E*^ drug candidates. The proposed therapeutics, engineered to incorporate constrained sp^3^-hybridized aliphatic rings and a sulfoximine bioisostere, demonstrated thermodynamically superior docking scores within the mutant catalytic cleft compared to dabrafenib. Lastly, a homology analysis determined that *Mus musculus* is a suitable model organism for future preclinical studies and confirmed crucial structural selectivity against microbial off-target kinases.

## INTRODUCTION

Melanoma is the third most prevalent cutaneous malignancy, posing a significant public health burden due to its aggressive clinical course and high metastatic potential. In contrast to nonmelanoma skin cancers, melanoma frequently metastasizes to the lungs, liver, bones, and central nervous system (CNS). With a five-year survival rate of 32% for stage IV metastatic melanoma as of 2024, there remains a critical need for advanced therapeutic interventions.^1^ Recently, the therapeutic paradigm has transitioned from traditional, low-efficacy cytotoxic chemotherapies to targeted agents focusing on the mitogen-activated protein kinase (MAPK) signaling cascade, a fundamental regulator of cellular proliferation.^1^ Targeting the MAPK is paramount due to its central role in driving unregulated cellular replication in metastatic disease.

### Molecular Mechanisms and BRAF Mutations

Molecularly, the MAPK/ERK pathway is triggered by extracellular stimuli that activate the small GTPase RAS. Upon binding GTP, RAS recruits RAF isoforms to the plasma membrane, promoting dimerization. In wild-type B-RAF kinases, dimerization is essential for catalyzing conformational transition from the inactive DFG-out state to the active DFG-in state through the extension of the activation loop, which allows the inward rotation of the αC -helix.^2,3^ During this transition, the aspartate residue of the DFG motif rotates toward the active site, facilitating ATP binding. The resulting active complex phosphorylates the downstream kinases MEK 1/2 and ERK 1/2, ultimately upregulating genes associated with cell survival.^2,3^

This physiological cascade is aberrantly activated by BRAF mutations in 40-50% of cutaneous melanomas. The most prevalent mutation is the *BRAF*^*V600E*^ variant, characterized by the substitution of valine with glutamic acid at position 600.^1,3,4^ This mutation destabilizes native hydrophobic interactions while concurrently stabilizing the active DFG-in state via the formation of a permanent salt bridge between Glu600 and Lys507.^3^

### The Structural Paradigm of Dimerization

The functional implications of this structural shift have been the subject of recent scientific reevaluation. Early models proposed that the stabilization of the DFG-in conformation allowed *BRAF*^*V600E*^ to participate exclusively in monomeric singling, effectively operating independently of upstream growth factor activation. However, contemporary structural studies have demonstrated that, much like wild-type BRAF, the mutant kinase relies heavily on heterodimerization to effectively phosphorylate MEK. This updated consensus is supported by evidence that *BRAF*^*V600E*^ possesses a physically expanded dimerization interface, readily forms dimers when expressed *in vivo*, and exhibits a severely reduced ability to induce MEK or ERK phosphorylation when paired with a kinase-dead receiver mutant incapable of binding MEK.^3^

Collectively, these findings indicate that while the *BRAF*^*V600E*^ mutation structurally locks the kinase in the DFG-in conformation, this stabilization does not bypass the need for dimerization. Rather, it allows the mutant kinase to spontaneously and aggressively assemble into dimers without requiring RAS-mediated membrane recruitment. This structural override serves as the primary mechanism for autoinhibition-resistant signaling and the uncontrollable cellular proliferation characteristic of metastatic melanoma.^3^

### Inhibitor Design and Structural Biology

Because BRAF mutations are primary oncogenic drivers in melanomagenesis, the rational design of therapeutic BRAF inhibitors to abrogate this aberrant signaling cascade has become a cornerstone in managing metastatic melanoma.^1^ These inhibitors demonstrate high clinical efficacy due to their preferential binding affinity for the mutated monomer relative to the physiologically active wild-type dimer.^5^ Binding predominantly occurs within the ATP-binding pocket, situated at the interface of N- and C-terminal lobes.^6^ Most of these inhibitors anchor to the kinase via hydrogen-bonding interactions with the hinge region backbone, notably at the Cys532 residue.^7^ Adjacent to the hinge region lies the hydrophobic adenine pocket, where the core scaffold of the inhibitor mimics the natural binding of ATP.^6^ Consequently, the mutant protein is stabilized in either an active (Type I) or inactive (Type II) conformation.^8^ Four prominent BRAF inhibitors — sorafenib, vemurafenib, dabrafenib, and encorafenib – exhibit well-characterized structural biology and clinical utility. The molecular structures of these therapeutic agents are illustrated in Figure 1.

**Figure 1.**
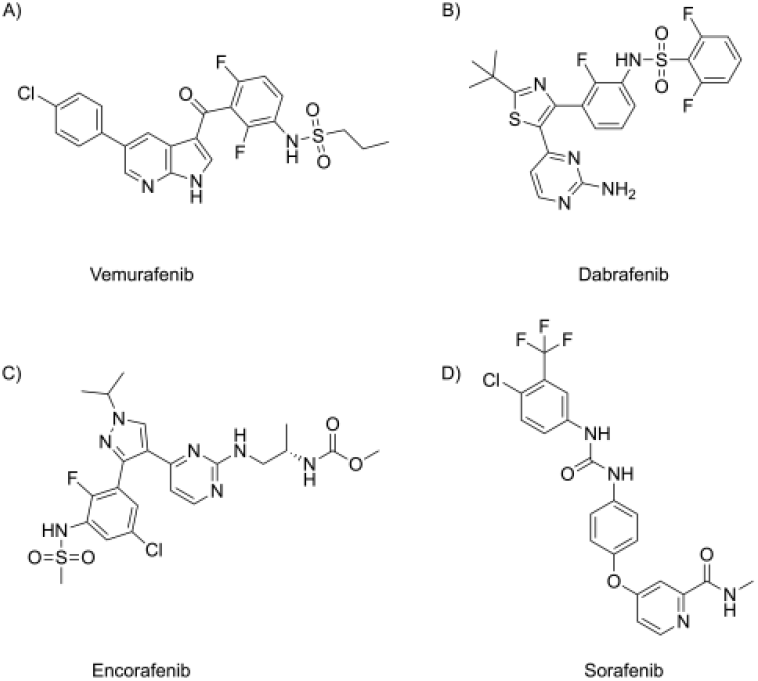
The molecular structures of first-generation BRAF kinase inhibitors: (A) vemurafenib, (B) dabrafenib, (C) encorafenib, and (D) sorafenib.

Despite their clinical success in the treatment of metastatic melanoma, the complex toxicological profiles of first-generation BRAF inhibitors warrant further structural optimization. A major clinical limitation of these agents is the rapid onset of acquired therapeutic resistance and paradoxical activation of the MAPK pathway in wild-type cells.^9^ Paradoxical activation occurs when sub-saturating drug concentrations stabilize a single protomer within a wild-type RAF dimer.^9^

Because RAF kinases function as an asymmetric pair, the drug-bound protomer is forced into an active αC-helix IN conformation, adopting the role of an allosteric activator.^10,11^ This activator structurally transactivates the drug-free partner kinase (the receiver), driving unregulated MEK phosphorylation and frequently fueling the development of secondary malignancies. ^3,11^

To circumvent this liability, recent developmental efforts have focused on “paradox-breaking” inhibitors designed to disrupt the enlarged *BRAF*^*V600E*^ dimer interface by targeting specific regulatory residues, such as Leu505 near the αC-helix.^5,12^ While this targeted approach successfully impairs dimerization, the current study hypothesizes that it creates a problematic thermodynamic reliance on narrow, highly localized “keystone” interactions. Because the inherent genomic instability of the melanoma creates a high probability that spontaneous point mutations will arise at these isolated nodes^13^, this hypothesized evolutionary vulnerability poses a significant hurdle to the clinical future of current paradox-breaking drug candidates, threatening to render them ineffective over time.

Proposing an alternative way to overcome this acquired resistance, the current study reconceptualizes the challenge of paradoxical activation by recognizing its inherent connection to the architectural limitations of early kinase inhibitors, which relied heavily on planar, aromatic, and achiral “privileged structures,” such as the versatile thiazole ring found in dabrafenib. While the flat, hinge-binding scaffolds of first-generation BRAF inhibitors provide robust target affinity, they lack the three-dimensional complexity required to avoid off-target interactions across the highly conserved kinase interactome. In recent years, the medicinal chemistry paradigm of “escaping from flatland” — incorporating complex, sp^3^-hybridized aliphatic systems and rigid stereocenters to maximize three-dimensional topological complementarity — has emerged as a validated strategy to combat this challenge.^14,15^

However, the current study argues that simply adopting this three-dimensional complexity is insufficient if these moieties remain structurally constrained to highly specific, mutable keystone residues. To establish a truly resilient steric blockade, an inhibitor must utilize this extended sp^3^ -rich bulk to create a distributed binding architecture spanning from the hinge region to the deep catalytic loop. This decentralized interaction network ensures that a single, random mutation is thermodynamically insufficient to compromise overall binding enthalpy.

Leveraging *in silico* computational methodologies, the current study sought to optimize the dabrafenib scaffold by systematically increasing its three-dimensional chemical complexity. Through the rational introduction of bulky sp^3^-hybridized ring systems, this study aimed to optimize dabrafenib by introducing broad mutation-resistant steric hindrance at the dimer interface (near αC-helix) to prevent off-target effects while also improving the *in-silico* binding affinity for the mutated protein. This rational design strategy yielded two novel compounds demonstrating enhanced computational binding affinity for the catalytically active *BRAF*^*V600E*^ conformation, engineered specifically for robust mutational resilience and the structural blockade of kinase dimerization.

## METHODS

### Protein Preparation and Active Site Definition

Three BRAF kinase crystal structures were retrieved from the Protein Data Bank (PDB).^16^ PDB IDs 4XV2^17^ and 3OG7^18^ represent the catalytically active *BRAF*^*V600E*^ mutant in the DFG-in conformation, co-crystallized with dabrafenib and vemurafenib, respectively. As depicted in Figure 2, the 4XV2-dabrafenib complex is characterized by the inward orientation of Asp594, diagnostic of the active DFG-in state.^3^ PDB ID 1UWH^19^ represents the wild-type kinase in the inactive DFG-out conformation, co-crystallized with sorafenib. These structures were selected to enable comparative binding affinity evaluations of DFG-in and DFG-out selecting inhibitors.^20^

**Figure 2.**
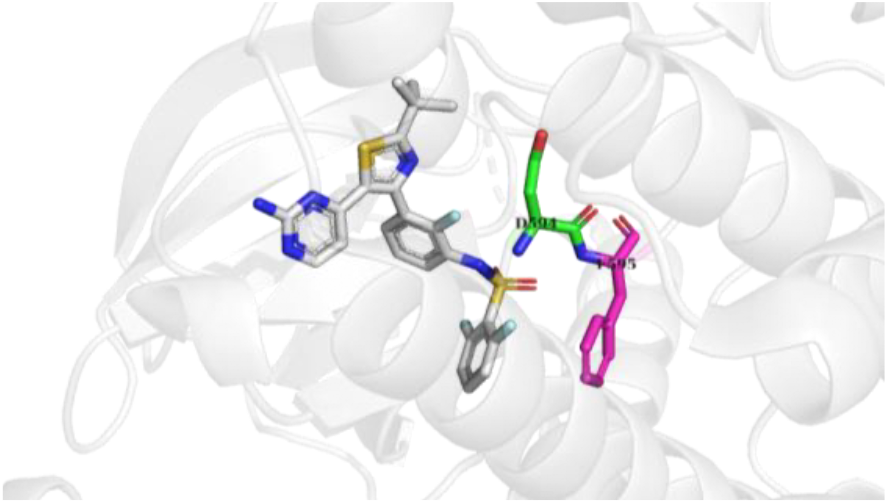
The DFG-in 4XV2 protein crystal structure in complex with dabrafenib

**Figure 3.**
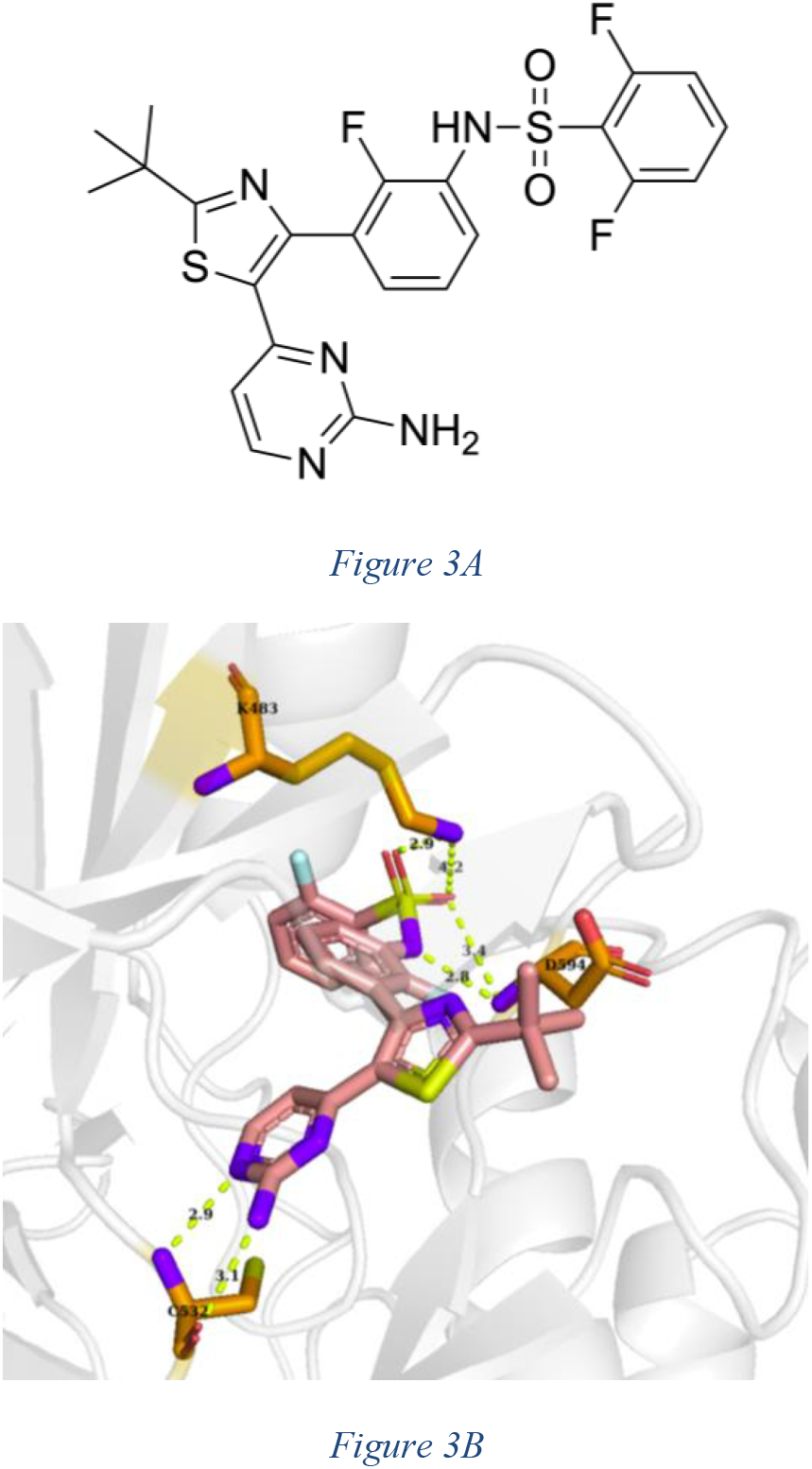
Binding profile of dabrafenib within the BRAF active site. (A) 2D chemical structure of dabrafenib. (B) Notable molecular interactions between the inhibitor and amino acid residues within the catalytic pocket.

Receptor active sites were defined utilizing the site detection and molecular probe algorithms within MakeReceptor (v. 4.1.1.0; OpenEye Scientific Software).^21^ Three-dimensional structures of control inhibitors (dabrafenib, encorafenib, sorafenib, and vemurafenib) were built in GaussView and subjected to AM1 semi-empirical geometry optimization using Gaussian 09.^22^ Ligand conformational ensembles were generated via OMEGA^23^ and subsequently docked into the respective receptor grids using FRED^24^ to establish baseline binding affinities and validate docking parameters.

### *De Novo* Design of Compound I

Compound I was identified via *de novo* design utilizing vBROOD (OpenEye).^25^ Two successive iterations of computational bioisosteric replacement were performed on the dabrafenib scaffold to generate a library of structural analogs. The resulting bioisosteres were evaluated via the validated FRED^24^ docking protocol, identifying Compound I as the top-scoring candidate.

### Structure-Based Lead Optimization (Compound II)

Compound II was derived via rational, structure-based lead optimization of Compound I. The predicted binding pose of the Compound I-4XV2 complex was visually analyzed in PyMOL^26^ to identify suboptimal contacts and unoccupied volume within the ATP-binding cleft. Targeted structural modifications designed to optimize local inhibitor interactions were constructed in GaussView, followed by AM1 geometry optimization in Gaussian 09.^24^ The refined derivatives were subjected to the established FRED^24^ docking protocol, yielding the fully optimized lead, Compound II.

### *In Silico* ADMET and Physiochemical Profiling

To evaluate pharmacokinetic viability, the control ligands, novel compounds (I and II), and intermediate derivatives were subjected to in silico physiochemical profiling using FILTER (OpenEye).^27^ Candidates failing to satisfy Lipinski’s Rule of Five (MW ≤ 500 Da, logP ≤ 5, Lipinski H-bond donors ≤ 5, Lipinski H-bond acceptors ≤ 10) or containing known toxicophores, aggregators, or reactive moieties (e.g., saponins)^28^ were systematically excluded. The resulting ADMET-compliant compounds were visualized and compiled into a refined target library utilizing VIDA.^29^

### Assessment of Off-Target Interactions

To evaluate cross-species selectivity and potential off-target liabilities, an *in silico* homology analysis was conducted utilizing NCBI Protein BLAST (blastp) against the human BRAF sequence (PDB ID 4XV2).^17^ The search was specifically directed against a commensal gut microbe (*Akkermansia muciniphila*)^31^, an opportunistic pathogen (*Staphylococcus aureus*)^32^, and a standard pre-clinical mammalian model (*Mus musculus*)^33^. Pairwise sequence alignments were generated via the NCBI Multiple Sequence Alignment Viewer (v. 1.26.0)^30^ to assess the evolutionary conservation of critical ATP-binding pocket residues: the hinge-region anchor (Cys532), the catalytic lysine (Lys483), and the DFG motif aspartate (Asp594). The resulting structural conservation data were utilized to model comparative binding affinities, predict potential off-target metabolic disruptions within the microbial species, and validate the translatability of the murine model for subsequent *in vivo* preclinical evaluations.

## RESULTS AND DISCUSSION

### Physicochemical Characterization of Established B-RAF Inhibitors

When subjected to an *in silico* ADMET-related property screen, as seen in Table I, sorafenib and vemurafenib fall within the traditional Lipinski Rule of Five limits for molecular weight (464.82 g/mol and 489.92 g/mol, respectively), though vemurafenib exhibits a slightly elevated lipophilicity with a xLogP of 5.44. Dabrafenib and encorafenib possess higher molecular weights (519.56 g/mol and 540.01 g/mol, respectively) that marginally exceed the 500 g/mol threshold. Encorafenib is the only compound in the dataset containing a chiral center and exhibits the highest number of hydrogen bond acceptors (11). Despite the slight molecular weight deviations for dabrafenib and encorafenib and encorafenib’s two Lipinski violations, all four compounds demonstrate viable pharmacokinetic profiles for kinase inhibition, with dabrafenib maintaining a favorable lipophilicity (xLogP = 4.16) and hydrogen bonding profile (HBD = 2, HBA = 7).

### Binding Affinity Analysis of Established B-RAF Inhibitors

Molecular docking simulations were executed to quantify the binding affinities of the control inhibitors across two catalytically active *BRAF*^*V600E*^ mutant crystal structures (PDB IDs: 4XV2, 3OG7)^17,18^ and one inactive wild-type structure (PDB ID 1UWH)^19^, utilizing the Chemgauss3^24^ scoring function (Table 2). A pronounced divergence in binding modalities was observed between the inhibitor classes when evaluated against mutant versus wild-type targets. These kinase inhibitors are structurally categorized into three classifications based on their selected protein conformations. The DFG-in state is characterized by the inward orientation of the Asp594 residue and the inward rotation of the N-terminal domain’s αC-helix toward the active site cleft. Accordingly, Type I inhibitors stabilize the DFG-in/aC-in conformation, Type II inhibitors stabilize the DFG-out/aC-out conformation, and Type IS inhibitors selectively stabilize the DFG-in/aC-out conformation. ^9,34^

**Table 1.** ADMET-related Properties of Known B-RAF Kinase Inhibitors.

| Compound | Chiral Center | Lipinski i HBD<sup>a</sup> | Lipinski i HBA<sup>b</sup> | MW<sup>c</sup> (g/mol) | xLog P |
| --- | --- | --- | --- | --- | --- |
| Dabrafenib | 0 | 2 | 7 | 519.56 | 4.16 |
| Encorafenib | 1 | 3 | 11 | 540.01 | 2.09 |
| Sorafenib | 0 | 3 | 7 | 464.82 | 3.73 |
| Vemurafenib | 0 | 2 | 6 | 489.92 | 5.44 |

<sup>a</sup>HBD = hydrogen bond donor, <sup>b</sup>HBA = hydrogen bond acceptor, <sup>c</sup>MW = molecular weight

**Table 2.** Molecular Docking Scores (Chemgauss3) ^24^ of B-RAF Inhibitors Across Mutant and Wild-Type Protein Crystal Structures.

| Compound<br>(Ligand) | Inhibitor<br>Type<sup>9,35</sup> | 4XV2<sup>17</sup><br>Score | 3OG7<sup>18</sup><br>Score | 1UWH<sup>19</sup><br>Score |
| --- | --- | --- | --- | --- |
|  |  | **V600E**<br>**Active** | **V600E**<br>**Active** | **Wild Type** |
| Sorafenib | II | -7.94 | -11.18 | -17.93 |
| Vemurafenib | IS<sup>d</sup> | -13.68 | -17.67 | -18.30 |
| Dabrafenib | IS | -10.60 | -11.27 | -7.98 |
| Encorafenib | IS | -7.81 | -9.36 | -7.00 |

<sup>d</sup>IS = type 1.5 inhibitors occupy the ATP site under DFG-in and αC-helix-out condition<sup>9,34</sup>

Sorafenib (Type II)^35^ and vemurafenib (Type IS)^9^ demonstrated robust overall binding affinities but exhibited a marked preferential binding for the wild-type 1UWH^19^ structure. Vemurafenib yielded Chemgauss3^24^ scores of -13.68 and -17.67 in the mutant complexes, yet achieved a superior score of -18.30 in the wild-type model. Similarly, the Type II^35^ inhibitor sorafenib recorded a highly favorable score of -17.93 in the wild-type structure compared to significantly attenuated affinities (-7.94 and -11.18) in the *BRAF*^*V600E*^ mutants, which is consistent with its established mechanism of action. Conversely, the Type IS^9^ inhibitors dabrafenib and encorafenib demonstrated pronounced selectivity for the *BRAF*^*V600E*^ mutant structures over the wild-type kinase. Dabrafenib exhibited binding scores of -10.60 (4XV2)^17^ and - 11.27 (3OG7)^18^ within the mutant active sites, contrasted by a significantly reduced affinity (-7.98) for the 1UWH^19^ wild-type. Encorafenib displayed a structurally analogous trend, albeit with generally attenuated overall binding affinities across the evaluated receptor ensemble.

### Rationale for the Selection of Dabrafenib for Lead Optimization

Based on this comparative analysis of physicochemical parameters and *in silico* docking affinities, dabrafenib was identified as the optimal structural scaffold for subsequent lead optimization.

The primary justification for this selection is its established structural selectivity. While vemurafenib yielded the greatest absolute binding scores, its failure to differentiate between the *BRAF*^*V600E*^ mutation and the wild-type kinase presents a substantial liability for off-target toxicity. In particular, the occurrence of this lack of specificity correlates clinically with vemurafenib’s higher propensity to induce paradoxical activation of the MAPK/ERK pathway, frequently driving the development of secondary malignancies in healthy cells.^36^ In contrast, dabrafenib successfully achieves the critical objective of mutant selectivity, preferentially binding to the active *BRAF*^*V600E*^ conformations, thereby mitigating the risk of paradoxical activation.

However, dabrafenib’s Chemgauss3^24^ scores in the mutant structures (-10.60 to -11.27) denote a moderate binding affinity, leaving a significant thermodynamic window for improvement. Consequently, dabrafenib serves as an ideal structural starting point: it inherently possesses the requisite spatial geometry to discriminate between mutant and wild-type BRAF yet affords ample opportunity for structural elaboration. Subsequent computational modifications will focus on the paradigm of “escaping from flatland^14^” by strategically introducing bulky sp^3^-hybridized ring systems and extending functional moieties deeper into the ATP-binding pocket. Ultimately, this increased three-dimensional complexity is intended to maximize Chemgauss3^24^ scores while simultaneously inducing broad steric hindrance at the dimer interface, yielding a compound optimized for both potency and mutational resilience.

### Structural Analysis of the Dabrafenib Binding Profile

Table 3 details the critical intermolecular contacts driving dabrafenib’s affinity for the *BRAF*^*V600E*^ active conformation. The baseline binding mode is defined by three primary anchor points within the ATP-binding pocket:

**Table 3.** Key Molecular Interactions and Distances Between Dabrafenib and BRAF (PDB:4XV2) ^17^.

| Residue Atom | Ligand Atom | Interaction Type | Distance (Å) |
| --- | --- | --- | --- |
| Cys532: N | Pyrimidine ring nitrogen | Hydrogen Bond | 2.9 |
| Cys532: O | Primary amine nitrogen | Hydrogen Bond | 3.1 |
| Asp594: OD1 | Sulfonamide nitrogen | Hydrogen Bond | 2.8 |
| Asp594: OD1 | Sulfonamide oxygen 2 | Hydrogen Bond | 3.4 |
| Lys 483: NZ | Sulfonamide oxygen 1 | Hydrogen Bond | 2.9 |
| Lys483: NZ | Sulfonamide oxygen 2 | Electrostatic Interaction | 4.2 |

### Hinge Region Anchoring (Cys532)

Dabrafenib establishes its core alignment by binding to the kinase hinge region, a hallmark interaction for ATP-competitive inhibitors.^6^ The pyrimidine ring nitrogen and the primary amine nitrogen form robust hydrogen bonds with the backbone nitrogen and oxygen of Cys532 at optimal distances of 2.9 Å and 3.1 Å, respectively.

### Engagement of the DFG Motif (Asp594)

The sulphonamide functional group acts as an effective dual hydrogen-bond donor and acceptor, heavily interacting with Asp594 (2.8 Å and 3.4 Å). Asp594 is the critical aspartate residue of the DFG motif located on the activation loop. The ability of dabrafenib to coordinate Asp594 stabilizes the kinase in the active “DFG-in” conformation to competitively inhibit ATP binding, an interaction characteristic of Type IS inhibitors.^9^

### Catalytic Lysine Interaction (Lys483)

The sulphonamide oxygens extend to engage Lys483, a conserved catalytic residue essential for ATP coordination.^8^ This engagement occurs via a strong 2.9 Å hydrogen bond and a supplementary 4.2 Å electrostatic interaction.

### Identifying Improvement Strategies and Structural Rationale

These molecular interactions validate the selection of dabrafenib as the primary computational scaffold for this study. The *in silico* data confirm that the core pyrimidine-sulfonamide backbone successfully targets the mechanistic vulnerabilities of the *BRAF*^*V600E*^ mutant. However, to systemically overcome the evolutionary and clinical limitations of first-generation inhibitors, the subsequent optimization campaign was directed by three distinct structural objectives:

1. **Disruption of the Dimerization Interface:** Consistent with a Type IS inhibitor^9^, dabrafenib exhibits strong selectivity for the DFG-in conformation, driven by its robust interaction with Asp594. In this active DFG-in conformation, the catalytic Lys483 residue is positioned in high proximity to the αC-helix, the primary structural component of the dimerization interface.^8,^ Consequently, optimizing and strengthening interactions with Lys483 provides a structural vector to project steric bulk toward the αC-helix, effectively disrupting the dimerization interface required to drive paradoxical activation in healthy cells.^2,3,8^
2. **Mitigation of Acquired Resistance:** Because Asp594 is prone to critical oncogenic missense mutations — such as the BRAF^D594G^ class 3 variant found frequently in lung adenocarcinomas^37^ — kinase inhibitors that invest a disproportionate ratio of their thermodynamic binding energy into this single coordinate are highly vulnerable to evasion. Crucially, while the D594G substitution renders the BRAF protomer catalytically “kinase-dead,” it paradoxically drives oncogenesis through a kinase-independent mechanism; the structural alteration orients the αC-helix into the active IN position, drastically increasing its dimerization potential and hyperactivating the MAPK pathway via transactivation of wild-type CRAF.^37^ This structural liability necessitates a thermodynamic shift in lead optimization, redirecting binding reliance away from the mutable Asp594 residue and toward enhanced, decentralized van der Waals networks deeper within the conserved binding cleft.
3. **Elimination of Photolability:** Recent pharmacokinetic profiling highlights a significant chemical liability in the baseline scaffold: dabrafenib is highly photolabile, undergoing rapid photoinduced degradation and photocyclization upon exposure to ultraviolet or ambient daylight in solution.^38^ This necessitates the targeted bioisosteric replacement of the planar, photolabile thiazole and difluorophenyl systems. To effectively address these three structural and clinical liabilities, the current study pursues strategic bioisosteric replacements of the planar, photolabile moieties. By substituting these two-dimensional systems with bulkier, sp^3^-hybridized aliphatic groups, the spatial footprint of the molecule is fundamentally expanded. This topological modification is explicitly designed to drive functional groups deeper into the lipophilic back pocket, intentionally shifting the thermodynamic reliance away from the known mutable active site residue, Asp594^38^, favoring a distributed, high-enthalpy binding network. By diverting a greater ratio of binding energy toward robust interactions, Lys483 and the deeper binding cleft, this extended three-dimensional bulk is projected directly toward the αC-helix interface. Ultimately, this structural evolution is hypothesized to establish a rigid steric blockade, simultaneously improving the inhibitor’s mutational resilience while combating the dimerization-driven paradoxical activation characteristic of first-generation therapeutics.^2,3,8^

### Rational Optimization and Binding Profile of Compound I

Guided by the structural liabilities of the baseline scaffold, Compound I was rationally designed to optimize spatial complementarity within the lipophilic back pocket of *BRAF*^*V600E*^.To achieve this, the OpenEye vBROOD^25^ bioisostere replacement search was utilized to strategically substitute the planar thiazole and difluorophenyl moieties with bulky, sp^3^-hybridized aliphatic systems intended to project deeper into the catalytic cleft.

### Enhanced Molecular Interactions and Binding Mode (PDB: 4XV2) ^17^

Molecular docking analyses (Figure 4B) and intermolecular distance measurements (Table 4) confirm that Compound I successfully retains the critical anchor points of the parental scaffold while establishing a superior, more robust distributed binding network within the mutant catalytic pocket:

**Table 4.** Key Molecular Interactions and Distances Between Compound I and BRAF (PDB:4XV2) ^17^.

| Residue Atom | Ligand Atom | Interaction Type | Distance (Å) |
| --- | --- | --- | --- |
| Cys532: N | Pyrimidine ring nitrogen | Hydrogen Bond | 2.9 |
| Cys532: O | Primary amine nitrogen | Hydrogen Bond | 3.4 |
| Cys532: N | Primary amine nitrogen | Weak Hydrogen Bond | 3.7 |
| Asp594: OD1 | Sulfonamide oxygen 1 | Hydrogen Bond | 3.3 |
| Lys 483: NZ | Sulfonamide oxygen 1 | Hydrogen Bond | 3.0 |
| Lys483: NZ | Sulfonamide<br>oxygen 2 | Hydrogen<br>Bond | 3.5 |

**Figure 4.**
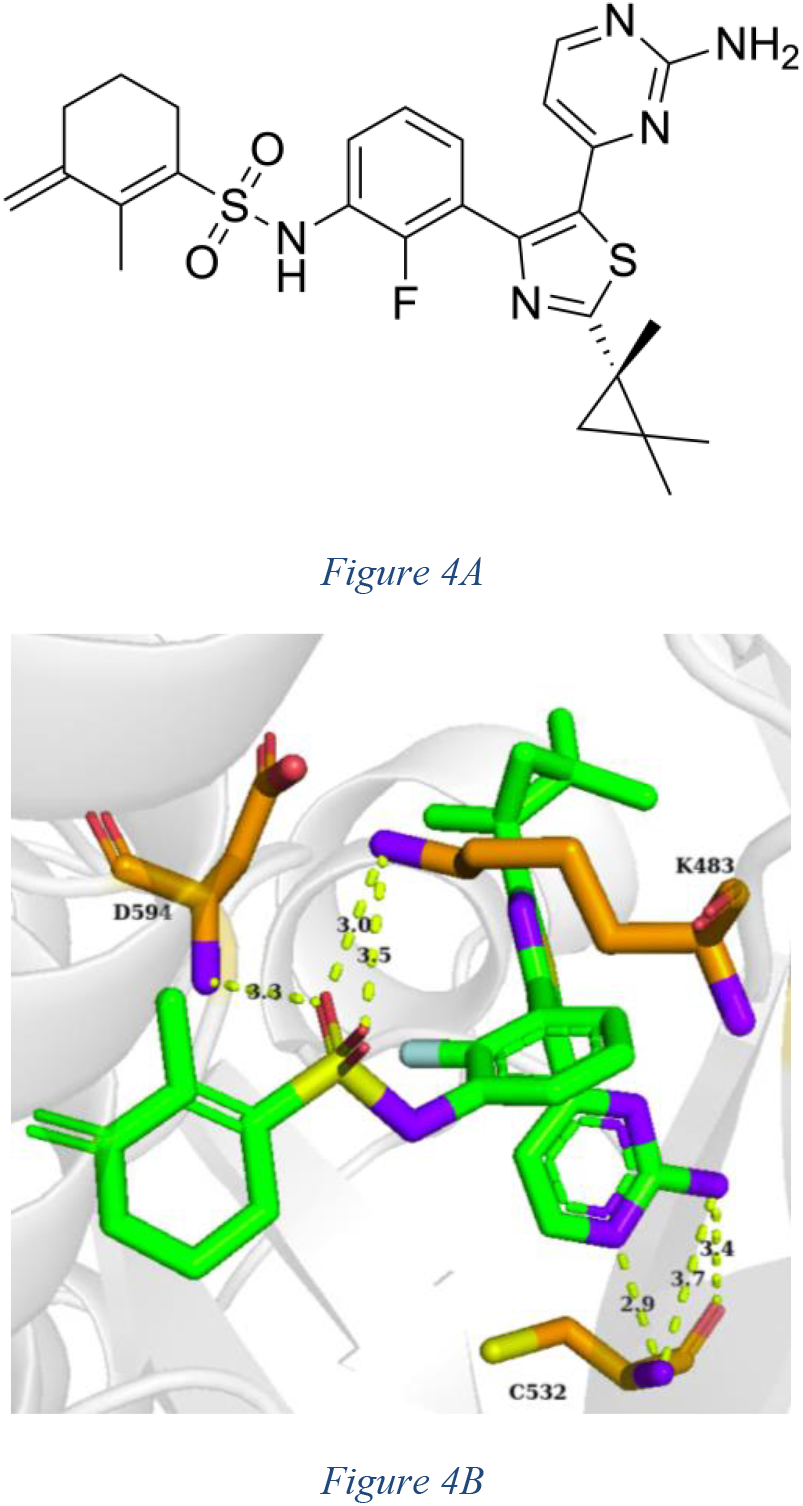
Binding profile of Compound I within the BRAF active site. (A) 2D chemical structure of Compound I. (B) Key molecular interactions between Compound I and amino acid residues in the catalytic pocket.

### Preservation of Hinge Anchoring

Compound I maintains absolute fidelity to the hinge region, with the pyrimidine ring nitrogen forming an optimal 2.9 Å hydrogen bond with the Cys532 backbone. The primary amine establishes a 3.4 Å hydrogen bond with the Cys532 carbonyl oxygen, supported by an auxiliary 3.7 Å interaction, securely locking the core heteroaromatic system in place.

### Novel Engagement Profile of the Catalytic Core (Lys483 and Asp594)

The most significant functional improvement over dabrafenib is observed deep within the catalytic cleft. While the baseline scaffold exhibited a single 2.9 Å hydrogen bond and a looser 4.2 Å electrostatic interaction with Lys483, Compound I achieves a significantly tighter, bidentate-like hydrogen-bonding network with the Lys483 NZ atom, measuring at 3.0 Å and 3.5 Å. The thermodynamic cost of this increased interaction with Lys483 is reflected in its altered interaction with Asp594; although Compound I maintains a strong 3.3 Å hydrogen bond with the OD1 atom of Asp594, the sulfonamide nitrogen is no longer at the required distance to form a second hydrogen bond.

This loss of the secondary Asp594 contact is a direct geometric consequence of incorporating the bulky, sp^3^- hybridized moieties. To accommodate this expanded spatial footprint within the binding cleft, the sulfonamide linker undergoes a subtle torsional shift, repositioning the heteroatoms. However, rather than representing a loss of affinity, this thermodynamic trade-off is highly advantageous. By sacrificing a secondary interaction with the mutable Asp594 active site residue in favor of a rigid, bidentate coordination with Lys483 — a strictly conserved residue essential for catalytic function^8,^ —Compound I shifts its energetic reliance toward a mutational “cold-spot” to prevent acquired resistance. Furthermore, anchoring the scaffold tighter against Lys483 effectively directs the newly introduced steric bulk toward the αC-helix, structurally aligning with the primary objective of disrupting the dimerization interface.

Further inspection of the consequences of this thermodynamic tradeoff was performed to evaluate the success of the current study’s second objective—to enhance van der Waals contacts deeper within the binding cleft. The emergence of novel non-electrostatic contact interactions in the Compound I-BRAF complex was identified using the FRED^24^-generated residue fingerprint (Table 5).

**Table 5.** FRED^24^-Identified Novel Contact Interactions Unique to the Compound I-BRAF Complex.

| Compound | Novel Contacts |
| --- | --- |
| Compound I | Asn580, Leu505, Phe468, Val 482 |

### Evaluation of the Distributed Binding Profile

The identification of these mutually exclusive contacts provides robust computational validation that the structural expansion successfully exploited the lipophilic recesses of the active site. The engagement of Phe468 is particularly signifcant, as this hydrophobic residue lines the ceiling and deeper regions of the ATP-binding cleft.^39^ Establishing van der Waals interactions with this specific amino acid confirms that the sp^3^-hybridized substitutions physically project deeper into the pocket than the planar dabrafenib scaffold. Additionally, the expanded spatial footprint allows the compound to pack efficiently against the αC-helix (Leu505) and the catalytic loop (Asn580).^12,40^ By distributing the binding energy across this broader network of hydrophobic contacts, rather than relying exclusively on a concentrated polar anchor at the DFG motif, Compound I establishes a substantial thermodynamic buffer. This comprehensive van der Waals optimization directly satisfies the second objective, establishing a distributed binding profile engineered to maintain target affinity even in the presence of localized conformational shifts or point mutations.

### Physicochemical and ADMET Profiling

The physicochemical impact of these structural modifications is detailed in Table 6. The integration of the bulkier aliphatic ring system appropriately increased the molecular weight to 539.69 g/mol and elevated the lipophilicity (xLogP = 4.88) compared to the dabrafenib scaffold (MW = 519.56 g/mol, xLogP = 4.16). This deliberate increase in xLogP fulfills the design objective of enhancing hydrophobic pocket engagement. Crucially, Compound I maintains an identical hydrogen bond donor (2) and acceptor (7) profile to dabrafenib, ensuring that the enhanced lipophilicity does not compromise essential solubility or permeability metrics. Furthermore, the introduction of a defined stereocenter imposes structural rigidity; this restricts the molecule’s rotational degrees of freedom, thereby entropically favoring the bioactive conformation necessary for *BRAF*^*V600E*^ selectivity and mutational resilience.^14^

**Table 6.** ADMET Properties of Compound 1.

| Compound | Chiral Center | Lipinski HBD | Lipinski HBA | Molecular Weight (g/mol) | xLogP |
| --- | --- | --- | --- | --- | --- |
| Compound I | 1 | 2 | 7 | 539.69 | 4.88 |

Concurrently, this structural evolution successfully fulfills the third optimization objective. By systematically replacing the highly conjugated, planar thiazole and difluorophenyl systems with saturated, sp^3^-hybridized aliphatic moieties, Compound I effectively eliminates the extensive pi-electron delocalization responsible for dabrafenib’s pronounced photolability. Consequently, this targeted bioisosteric switch neutralizes the primary chemical liability of the baseline scaffold while maintaining strict adherence to Lipinski’s Rule of Five, yielding a highly stable, pharmacokinetically viable lead compound.^38^

### Summary of Improvements

Compound I represents a significant structural evolution over dabrafenib. By overhauling the photolabile planar moieties with a bulky, chiral sp^3^-architecture, the molecule deeply exploits the lipophilic back pocket while tightening coordination at the catalytic Lys483. This enhanced thermodynamic grip, combined with entropically favorable rigidification, establishes a comprehensive steric blockade at the αC-helix. Ultimately, this structural optimization shows potential for the mutation-resistant, physical disruption of the dimerization interface.

### Rational Design and Sulfoximine Bioisosterism in Compound II

Compound II (Figure 5A) was engineered as a refined evolutionary step to optimize the thermodynamic profile of the scaffold and address residual physicochemical liabilities. While retaining the crucial sp^3^-hybridized steric bulk utilized in earlier iterations to disrupt the dimerization interface, the defining structural modification in Compound II is the bioisosteric replacement of the classical sulfonamide linkage with a sulfoximine moiety.

**Figure 5.**
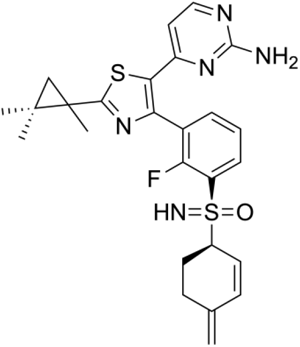
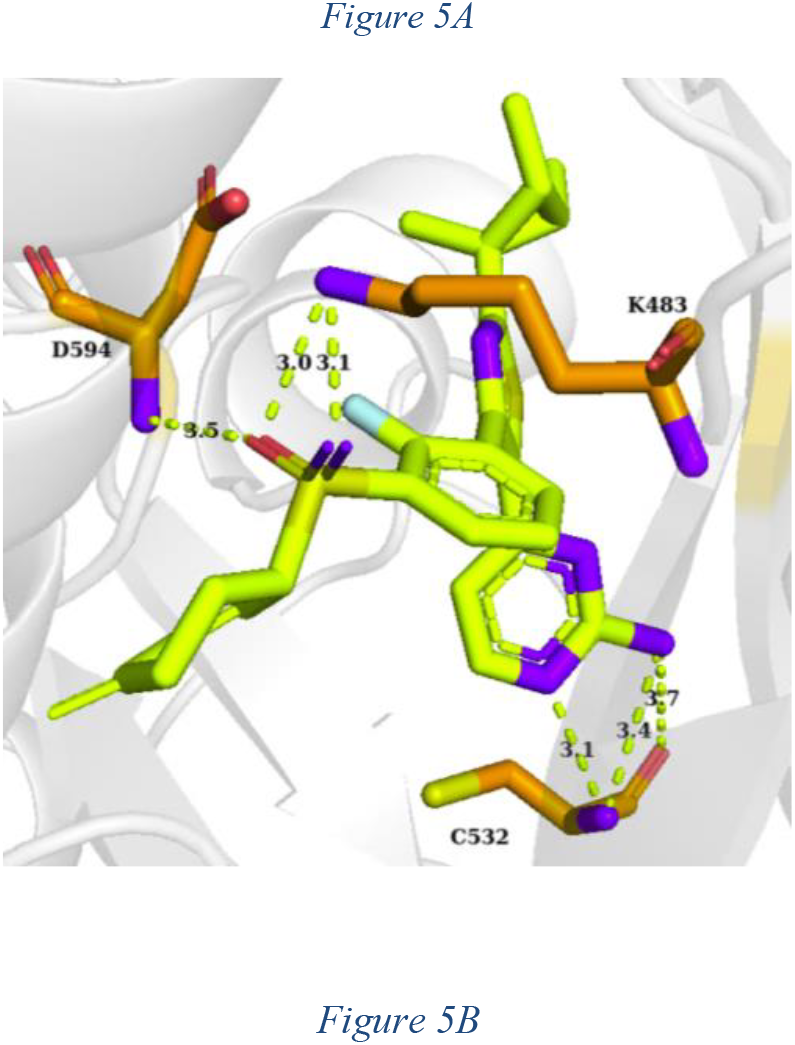
Binding profile of Compound II within the BRAF active site. (A) 2D chemical structure of Dabrafenib. (B) Key molecular interactions between Compound II and amino acid residues in the catalytic pocket.

Sulfoximines have emerged as highly privileged, three-dimensional bioisosteres in modern medicinal chemistry, offering distinct pharmacological advantages over sulfonamides.^41^ These advantages include enhanced aqueous solubility, remarkable metabolic resilience, and an avoidance of sulfonamide-associated cross-reactivities and toxicities.^41^ Furthermore, the tetrahedral geometry of the sulfoximine sulfur atom inherently introduces a stereogenic center.^42^ By adding this chiral center to the existing constrained scaffold, Compound II undergoes profound rigidification. This restriction entropically pre-organizes the molecule into the bioactive DFG-in conformation, maximizing target affinity while minimizing the conformational flexibility that often drives off-target kinome engagement.^43^

### Elevated Binding Network and Bidentate Coordination

Molecular docking against the *BRAF*^*V600E*^ active conformation (Figure 5B and Table 7) reveals that the sulfoximine substitution fundamentally elevates the binding network within the catalytic core, representing a clear structural advantage over Compound I.

**Table 7.**
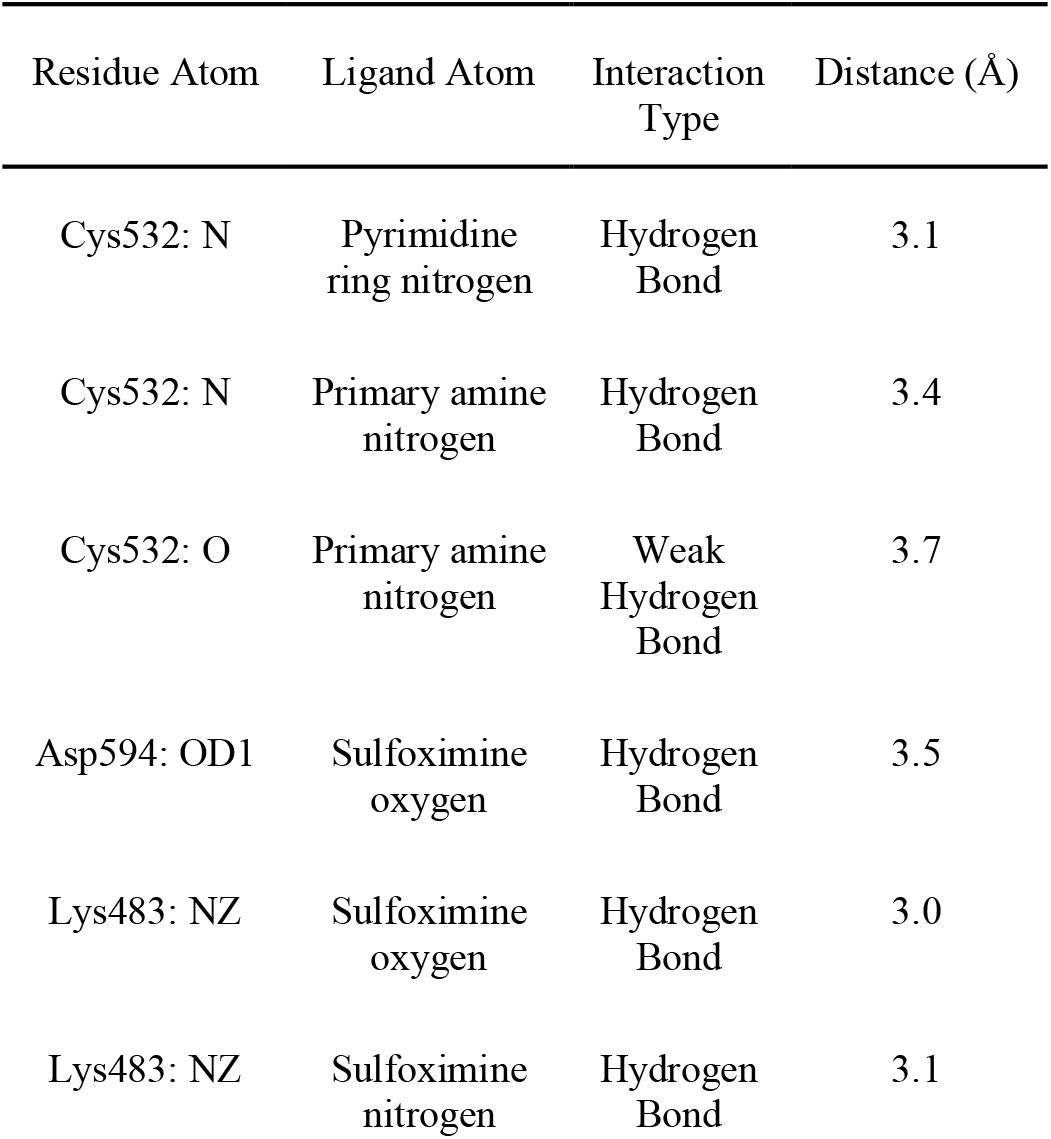
Key Interactions and Distances Between Compound II and BRAF (PDB:4XV2) ^17^.

Compound II maintains the indispensable hinge anchoring at Cys532 so that the core heteroaromatic system remains securely aligned. However, a significant thermodynamic improvement is observed at the critical interface between the catalytic Lys483 and the DFG motif (Asp594). In both dabrafenib and Compound I, target engagement in this region was mediated exclusively by sulfonamide oxygen atoms. While oxygens are effective hydrogen-bond acceptors, their high electronegativity causes them to hold their lone pairs tightly, limiting maximum bond enthalpy.^44^ The bioisosteric switch to a sulfoximine dynamically alters this electronic landscape by introducing a nitrogen atom into the coordination sphere. Because nitrogen is less electronegative and more polarizable than oxygen^44^, its electron density is more readily available to interact with the amino group (NZ) of Lys483. Consequently, Compound II exhibits lessened reliance on the known mutable Asp594 residue (distance extending to 3.5 Å), transferring that thermodynamic weight to an exceptionally tight, bidentate grip on the strictly conserved Lys483. The sulfoximine oxygen forms a strong 3.0 Å hydrogen bond, while the superior electron-donating capacity of the sulfoximine nitrogen drives a highly directional 3.1 Å hydrogen bond. This bidentate coordination firmly anchors the scaffold within the active site and accomplishes the structural objective of disrupting the dimerization interface.

Before discussing the broader clinical implications of the improvements introduced in the novel Compound II, it must be noted that while the compound fundamentally upgrades the interaction at the catalytic lysine, it successfully preserves the expanded hydrophobic footprint first established by Compound I. This includes the novel contact network within the extended pocket (Gly593, Ile513, Leu505, Phe595, and Thr529) identified via FRED docking (Table 8).

**Table 8.** FRED-Identified Novel Contact Interactions Unique to the Compound II-BRAF Complex.

| Compound | Novel Contacts |
| --- | --- |
| Compound II | Gly593, Ile513, Leu505, Phe595, Thr529, |

### Therapeutic Implications of Compound II: Overcoming Resistance and Toxicity

#### Mutational Resilience

The bidentate interaction with Lys483, facilitated by the valency of the sulfoximine nitrogen, significantly deepens the thermodynamic well of the binding event. This high binding affinity provides a robust energetic buffer against acquired drug resistance, enabling the inhibitor to maintain efficacy even if secondary mutations alter the conformational dynamics of the broader binding pocket.^3,9,11^

### Suppression of Paradoxical Activation

The structural rigidity imparted by the three chiral centers constrains the molecule’s spatial footprint. This rigidified geometry strongly favors the *BRAF*^*V600E*^ mutant conformation while creating massive steric hindrance at the αC-helix. This structural blockade physically prevents the drug-bound complex from functioning as an allosteric activator, thereby abrogating the heterodimerization interfaces required to transactivate wild-type receiver kinases in healthy cells, effectively mitigating the risk of secondary malignancies.^3,43,^

Table 9 summarizes the physicochemical profile of Compound II. While the foundational sp^3^ architecture of Compound I was largely conserved to maintain deep pocket engagement, Compound II incorporates a targeted structural truncation: the explicit removal of the methyl substituent from the aliphatic ring system. This deliberate demethylation, combined with the sulfoximine bioisosterism, strategically reduces the overall molecular weight to 509.66 g/mol and modulates the lipophilicity to a more favorable xLogP of 4.57. This structural refinement strikes a highly optimized pharmacokinetic balance. It successfully dials back the elevated lipophilicity of Compound I (xLogP = 4.88) to improve aqueous solubility, while remaining significantly more lipophilic than the dabrafenib baseline to ensure robust exploitation of the hydrophobic cleft. The essential hydrogen bond donor capacity (HBD = 2) is perfectly conserved. Concurrently, Compound II definitively fulfills the third optimization objective of this study. By replacing the highly conjugated, planar thiazole and difluorophenyl systems of earlier generations with a saturated sp^3^-architecture, and exchanging the sulfonamide for a metabolically robust sulfoximine, the primary chemical liability of photolability is completely eradicated.^38^ This ADMET profile confirms Compound II as a photostable, highly soluble, and optimally lipophilic lead compound that maximizes on-target potency while strictly adhering to Lipinski’s Rule of Five.

**Table 9.** ADMET Properties of Compound II.

| Compound | Chiral Center | Lipinski HBD | Lipinski HBA | Molecular Weight (g/mol) | xLogP |
| --- | --- | --- | --- | --- | --- |
| Compound II | 3 | 2 | 6 | 509.66 | 4.57 |

**Table 10.** Molecular Docking Scores (Chemgauss3) ^24^ of Dabrafenib, Compound I, and Compound II Across Mutant and Wild-Type Protein Crystal Structures.

| Compound | Known or Proposed | 4XV2<sup>17</sup> (V600E) | 3OG7<sup>18</sup> (V600E) | 1UWH<sup>19</sup> Score |
| --- | --- | --- | --- | --- |

| (Ligand) | Inhibitor Type | Active) | Active) | (WT) |
| --- | --- | --- | --- | --- |
| Dabrafenib | IS | -10.60 | -11.27 | -7.98 |
| Compound I | IS | -13.62 | -12.88 | -2.32 |
| Compound II | IS | -14.33 | -14.99 | -2.39 |

### Thermodynamic Validation and Mutant Selectivity via *In Silico* Scoring

The structural and electronic optimizations of Compound I and Compound II are quantitatively validated by their respective OpenEye Chemgauss3^24^ docking scores across both *BRAF*^*V600E*^ mutant (4XV2, 3OG7)^17,18^, and Wild-Type (1UWH)^19^crystal structures.

Against the mutant active conformations, the dabrafenib baseline exhibited binding scores of -10.60 and -11.27. Compound I demonstrated significantly improved scores (- 13.62 and -12.88), providing robust computational evidence that establishing a distributed van der Waals network effectively compensates for the partial loss of the DFG-motif hydrogen-bonding network (the Asp594 residue). Compound II represents the thermodynamic apex of this optimization campaign, achieving vastly superior mutant docking scores of -14.33 and -14.99. This quantitative progression definitively supports the hypothesis established in prior sections: severing reliance on the mutable Asp594 residue and redirecting the binding energy into a localized, bidentate coordination of Lys483 successfully generates a highly favorable, mutation-resilient complex.

### Elimination of Wild-Type Affinity and Paradoxical Activation

Crucially, the Chemgauss3^24^scores against the Wild-Type BRAF structure (1UWH)^19^ confirm that this thermodynamic shift confers an unprecedented degree of mutant selectivity. While dabrafenib exhibits notable off-target affinity for the Wild-Type kinase (-7.98) — the primary interaction that drives the clinical liability of paradoxical activation — the sp^3^-hybridized constraints of Compound I and Compound II severely penalize Wild-Type engagement, plummeting their respective scores to -2.32 and -2.39. This massive differential computationally validates the structural blockade discussed previously. By mathematically abolishing affinity for the Wild-Type kinase, Compound I and Compound II are rendered less prone to inducing the allosteric activator conformation in healthy cells.^2,3,38^ Ultimately, these superior docking scores confirm that “escaping from flatland^14^” not only yields a more potent complex against *BRAF*^*V600E*^, but also fundamentally engineers the selectivity required to eradicate paradoxical dimerization and off-target toxicity.

### Homolog Analysis

Beyond exhibiting superior binding affinity and mutational resilience within the human *BRAF*^*V600E*^ kinase, Compound I and Compound II show therapeutic utility further validated by a comprehensive *in silico* homolog analysis. This safety profiling serves to define the translatability of the novel compounds into pre-clinical animal models while simultaneously assessing the risk of off-target activity against critical components of the human commensal microbiome and common pathogenic flora.

### Pre-clinical Animal Model Analysis, *Mus musculus*, (ID: 10088

The serine/threonine protein kinase of *Mus musculus* was identified as a functional homolog of human BRAF (Sequence ID: AAA37320.1). High-resolution structural alignment of the kinase domain confirms near-total conservation of the catalytic machinery, including the DFG motif (Asp594), the catalytic loop (Lys483), and the hinge-region anchor (Cys532). Given that these residues comprise the essential hydrogen-bonding network prerequisite for ATP-competitive inhibition^3^, Compound I and Compound II are predicted to exhibit high-affinity binding to the murine homolog. This structural conservation identifies *Mus musculus* as a highly representative pre-clinical model, providing a robust system to evaluate the pharmacokinetic profile, therapeutic index, and systemic toxicity of the two novel compounds prior to human clinical translation.

### Commensal Microbe Safety Profile, *Akkermansia muciniphila* (id: 239935)

The serine/threonine protein kinase of the beneficial microbe *Akkermansia muciniphila* was identified as a homolog of the human BRAF protein kinase with a percent identity of 30.29%. The critical hinge-region residues, Cys532 and Lys483, are not conserved and are replaced by asparagine (Asn) residues in the microbial sequence. The active-site DFG motif (Asp594), however, is conserved between the two sequences, indicating identical structural machinery for ATP binding in this homolog.^3^ While this motif remains intact, Cys532 and Lys483 are essential for the binding of ATP-competitive BRAF kinase inhibitors.^3^ Therefore, Compound I and Compound II are not expected to bind with high affinity to the homologous bacterial protein. This suggests that the compounds will not inhibit the *A. muciniphila* kinase, allowing the microbe’s metabolic signaling pathways to remain functional without impacting its capacity for survival and cellular proliferation. *Akkermansia muciniphila* is a crucial component of the human gut microbiome, playing essential roles in combatting various metabolic and systemic conditions, including type 2 diabetes, inflammation, inflammatory bowel disease (IBD), and cardiovascular disease.^45^ Ultimately, this structural selectivity is highly promising, ensuring the targeted cancer therapy will not inadvertently decimate beneficial gut flora.

### Pathogenic Microbe Homolog Analysis, *Staphylococcus aureus*, (ID: 1280)

The serine/threonine protein kinase of the pathogenic microbe *Staphylococcus aureus* was identified as a homolog of the human BRAF protein kinase. Upon structural alignment, the active-site DFG motif (Asp594) and the catalytic loop (Lys483) are fully conserved between the two sequences, indicating identical structural machinery for ATP binding and facilitating active catalysis in this homolog. However, the critical hinge-region residue, Cys532, is not conserved and is replaced by a valine (Val) residue in the microbial sequence. While the core catalytic machinery remains intact, Cys532 is essential for providing the specific hydrogen-bonding network required to anchor ATP-competitive BRAF kinase inhibitors.^3^ The substitution with a bulky, hydrophobic valine introduces a steric and electrostatic barrier. Therefore, Compounds I and II are not expected to bind with high affinity to the homologous bacterial protein, allowing the pathogen’s metabolic signaling pathways to remain fully functional. Consequently, the drugs will not exhibit off-target antimicrobial activity against this *Staphylococcus* species. While this means the compounds cannot serve as dual-action therapeutics to concurrently treat a Staph infection, this high structural selectivity is ultimately a major advantage for patients being treated for metastatic melanoma, as it ensures the targeted cancer therapy will not inadvertently contribute to the development of widespread antibiotic resistance.^46^

## CONCLUSION

The continuous emergence of acquired drug resistance and paradoxical MAPK pathway activation necessitates the evolution of next-generation *BRAF*^*V600E*^ inhibitors. In this study, the Type IS inhibitor dabrafenib was computationally optimized to address its reliance on mutable active-site residues and its susceptibility to off-target wild-type dimerization. Through de novo VBROOD bioisosteric replacement, Compound I successfully incorporated a bulky, sp^3^-hybridized aliphatic architecture that extended the van der Waals interaction network deeper into the lipophilic catalytic cleft. Subsequent rational structural optimization yielded Compound II, which replaced the photolabile, planar sulfonamide with a rigid, three-dimensional sulfoximine bioisostere. *In silico* docking studies validated that Compound II achieves vastly superior thermodynamic binding affinities against the active *BRAF*^*V600E*^ mutant conformation (Chemgauss3 score: -14.99) compared to the dabrafenib baseline (-11.27). Crucially, this optimization successfully shifted the thermodynamic reliance from the mutable Asp594 residue to a highly directional, bidentate coordination with the strictly conserved Lys483. This anchoring firmly projects the molecule’s steric bulk toward the αC-helix, computationally severely penalizing wild-type engagement (Chemgauss3 score: -2.39) to establish the structural blockade required to abrogate paradoxical dimerization. Finally, a comprehensive homology analysis confirmed the translational viability of Compounds I and II. High structural conservation of the kinase domain verified *Mus musculus* as an appropriate preclinical model, while crucial deviations at the Cys532 and Lys483 hinge residues in *Akkermansia muciniphila* and *Staphylococcus aureus* predict excellent safety profiles against both commensal and pathogenic microbiomes. Ultimately, Compounds I and II represent highly optimized, ADMET-compliant, and mutation-resilient lead candidates that effectively navigate the complex structural biology of metastatic melanoma.

## Supporting information

File S1: BRAF Docking Reports

File S2: BLAST Alignments

File S3: Methodological Transparency

## ACKNOWLEDGMENTS

All informatics and molecular modeling, as well as modeling interpretation, were conducted by Z.H.Y. The manuscript was reviewed by E.R.M. and J.B.S., and Gemini 3.1 was utilized strictly for copyediting of the final draft.

This work was supported by the University of California, Davis, the National Science Foundation (award nos. # 2315767, 2118138, and 1827246), and BioMADE FA8650-21-2-5028. The content is solely the responsibility of the authors and does not necessarily represent the official views of the National Science Foundation, other sponsors, or UC Davis.

