## Supplementary material for "Computational Design of Two Novel BRAF V600E Inhibitors: Exploiting Sulfoximine Bioisosterism and Chiral Constraints to Evade Paradoxical Activation": File S1: BRAF Docking Reports

| | |
|---|---|
| Molecule Name | Vemurafenib_85 |
| Molecular Weight | 489.9 |
| XLogP | 5.4 |
| PSA | 91.9 |
| Heavy Atoms | 33 |
| Acceptor Count | 4 |
| Donor Count | 2 |
| Chelator Count | 1 |

**PDB ID: 4XV2**

### Total Score -13.68

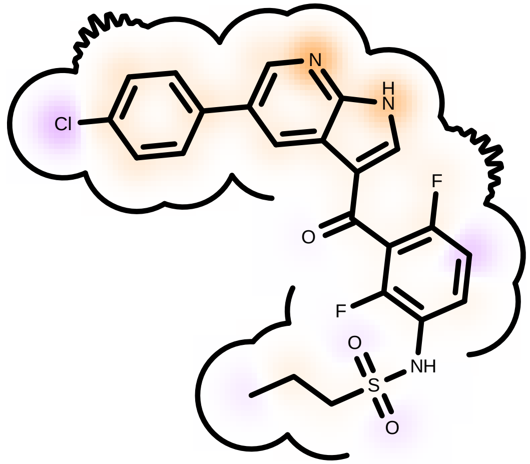

Score compared to other molecules

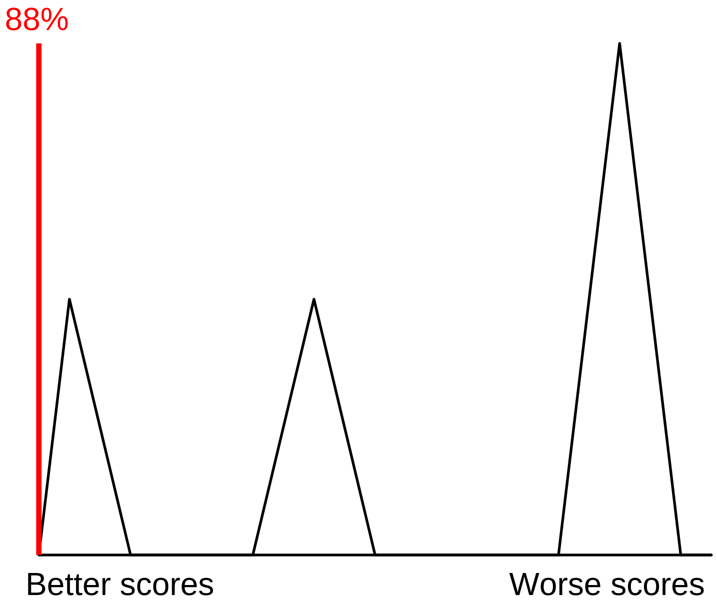

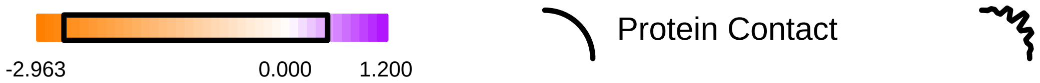

Residue Fingerprint

| | |
|---|---|
| ALA481A | ASN580A |
| ASP594A | CYS532A |
| GLN530A | GLU533A |
| GLY464A | GLY466A |
| GLY534A | GLY593A |
| ILE463A | ILE527A |
| LEU505A | LEU514A |
| LYS483A | PHE468A |
| PHE583A | PHE595A |
| SER465A | SER535A |
| SER536A | THR529A |
| TRP531A | VAL471A |

| | |
|---|---|
| Acceptor | Donor |
| Metal | Contact |

Shape -13.75

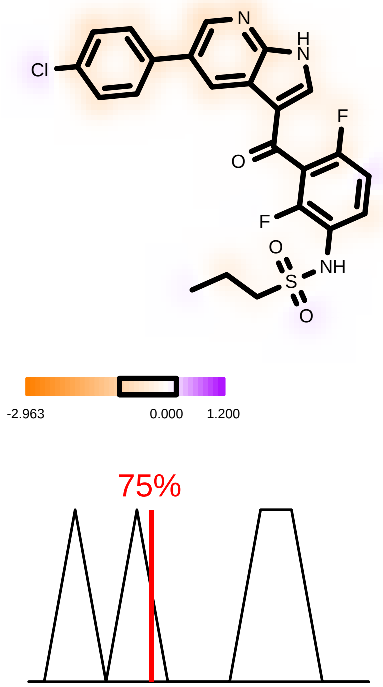

Hydrogen Bond -5.76

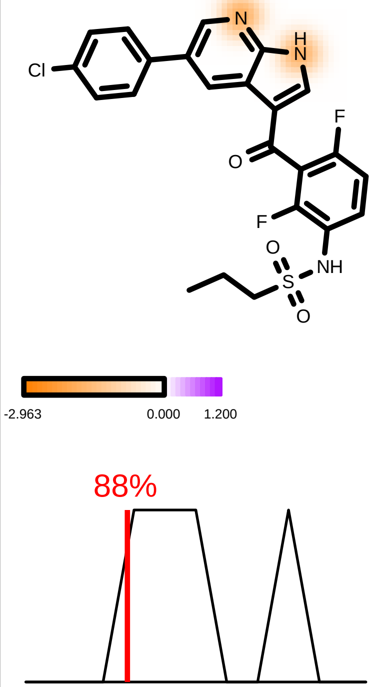

Protein Desolvation 3.01

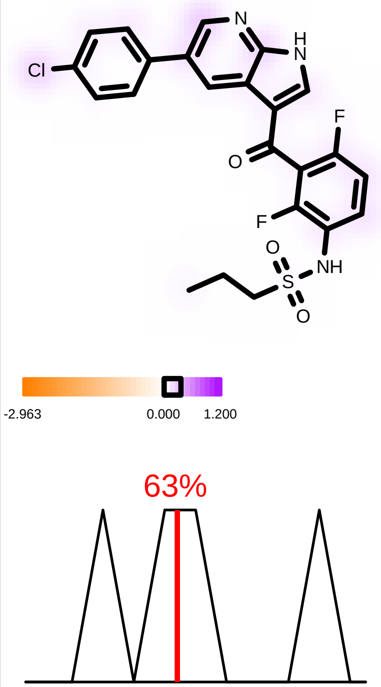

Ligand Desolvation 2.82

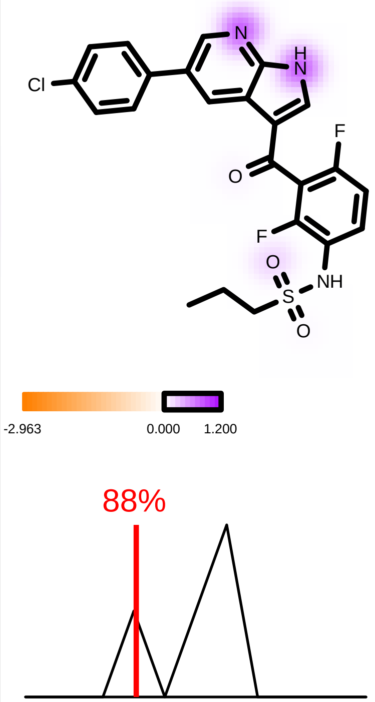

| | |
|---|---|
| Molecule Name | Molecule Name 139 |
| Molecular Weight | 519.6 |
| XLogP | 5.0 |
| PSA | 110.9 |
| Heavy Atoms | 35 |
| Acceptor Count | 6 |
| Donor Count | 2 |
| Chelator Count | 3 |

PDB ID: 4XV2

***(Dabrafenib)***

### Total Score -10.60

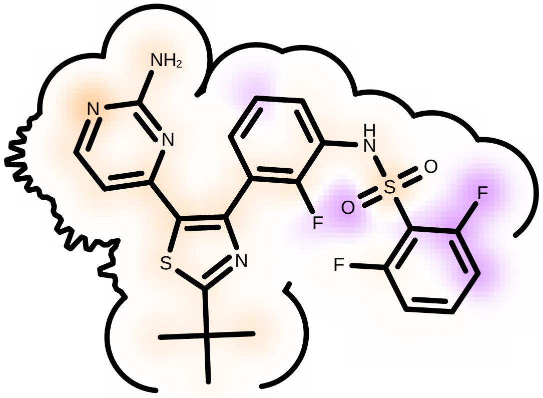

Score compared to other molecules

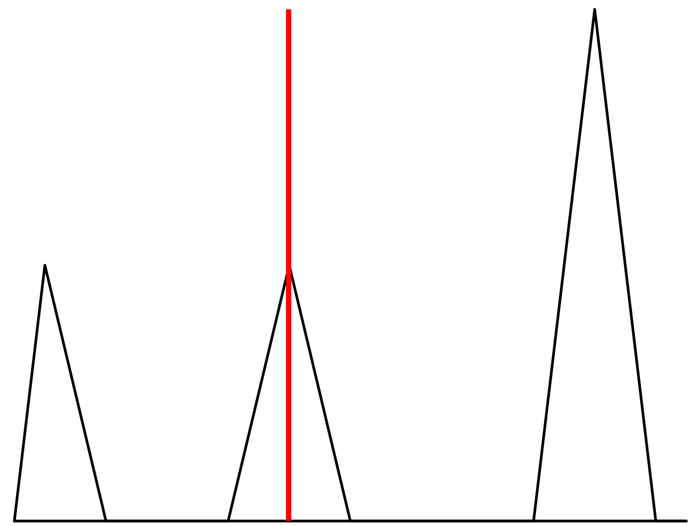

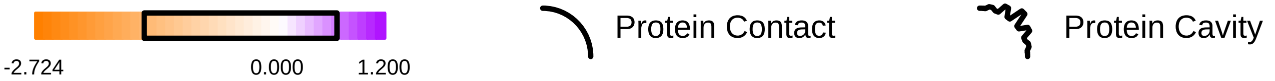

Residue Fingerprint

| | |
|---|---|
| ALA481A | ASN580A |
| ASP594A | CYS532A |
| GLN530A | GLU533A |
| GLY464A | GLY466A |
| GLY534A | GLY593A |
| ILE463A | ILE527A |
| LEU505A | LEU514A |
| LYS483A | PHE468A |
| PHE583A | PHE595A |
| SER465A | SER535A |
| SER536A | THR529A |
| TRP531A | VAL471A |

Acceptor Donor
Metal Contact

Shape -11.58

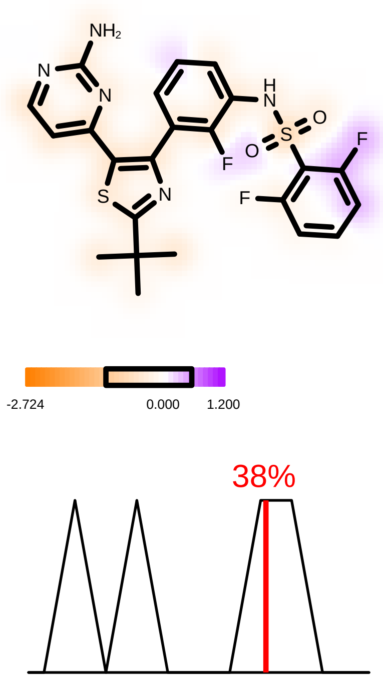

Hydrogen Bond -4.85

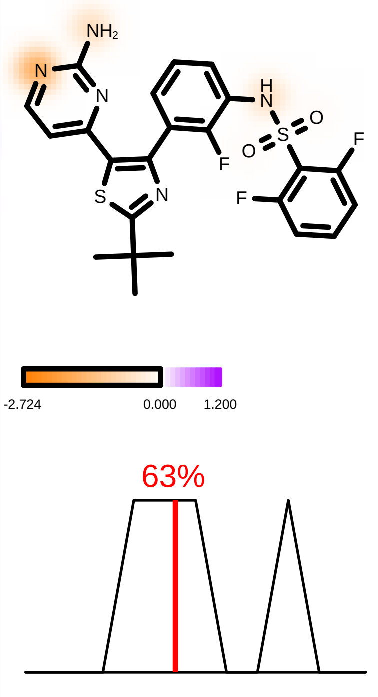

Protein Desolvation 1.83

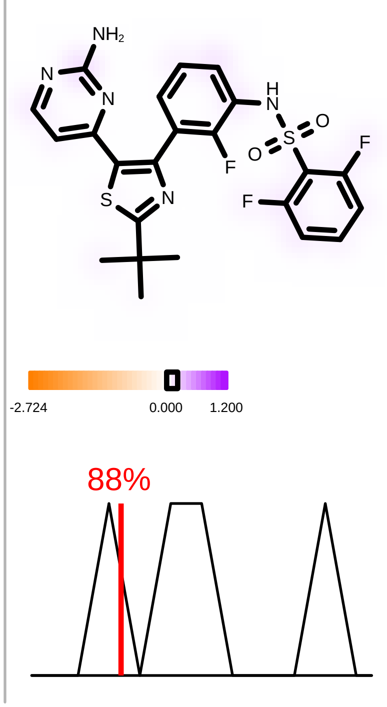

Ligand Desolvation 4.00

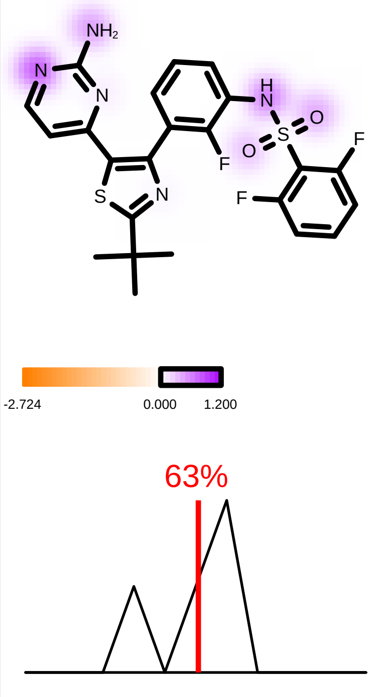

| | |
|---|---|
| Molecule Name | Sorafenib_110 |
| Molecular Weight | 464.8 |
| XLogP | 3.7 |
| PSA | 92.4 |
| Heavy Atoms | 32 |
| Acceptor Count | 4 |
| Donor Count | 3 |
| Chelator Count | 1 |

PDB ID: 4XV2

### Total Score -7.94

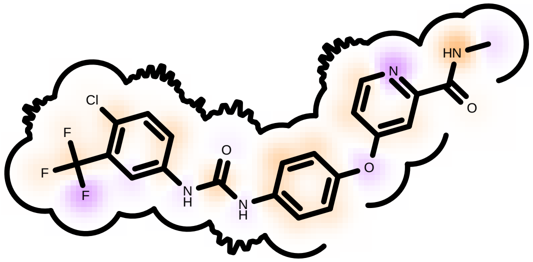

Score compared to other molecules

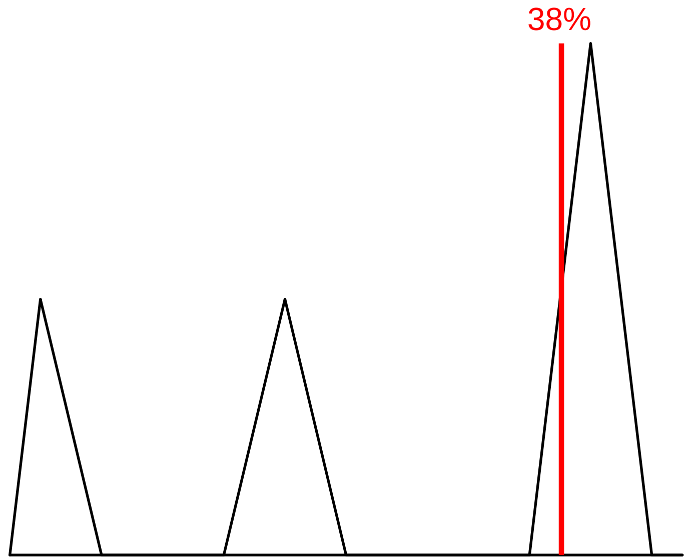

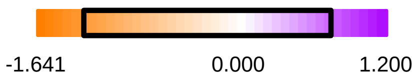

Protein Contact

Protein Cavity

Residue Fingerprint

| | |
|---|---|
| ALA481A | ASN580A |
| ASP594A | CYS532A |
| GLN530A | GLU533A |
| GLY464A | GLY466A |
| GLY534A | GLY593A |
| ILE463A | ILE527A |
| LEU505A | LEU514A |
| LYS483A | PHE468A |
| PHE583A | PHE595A |
| SER465A | SER535A |
| SER536A | THR529A |
| TRP531A | VAL471A |

Acceptor Donor
Metal Contact

Shape -15.23

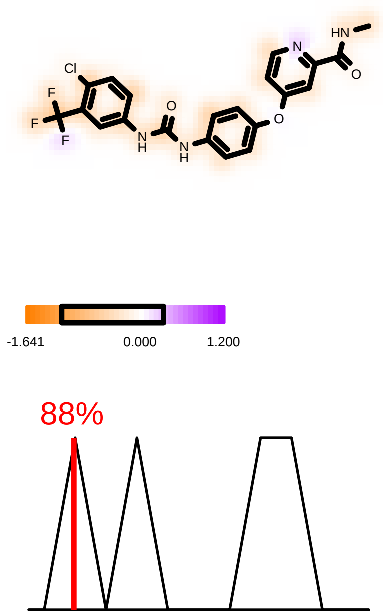

Hydrogen Bond -2.60

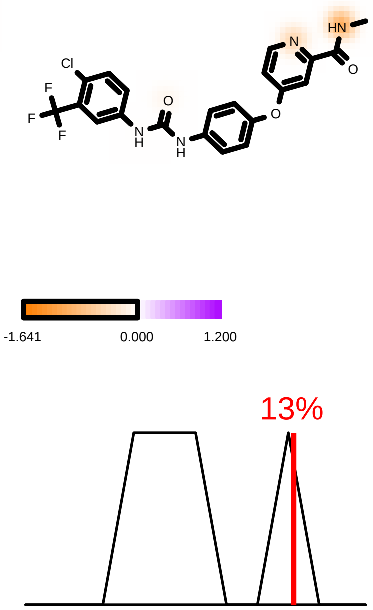

Protein Desolvation 5.44

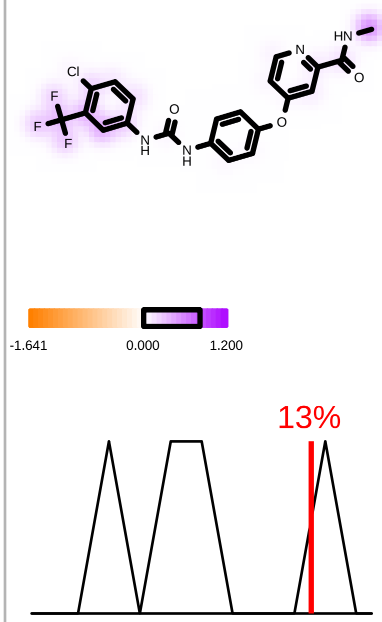

Ligand Desolvation 4.44

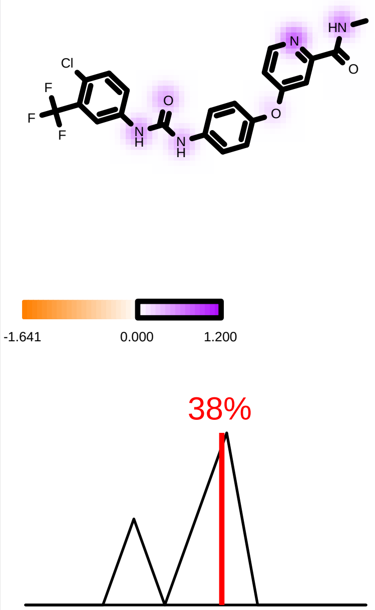

| | |
|---|---|
| Molecule Name | Molecule Name 155 |
| Molecular Weight | 540.0 |
| XLogP | 2.6 |
| PSA | 140.1 |
| Heavy Atoms | 36 |
| Acceptor Count | 6 |
| Donor Count | 3 |
| Chelator Count | 3 |

*(Encorafenib)*

**PDB ID: 4XV2**

### Total Score -7.81

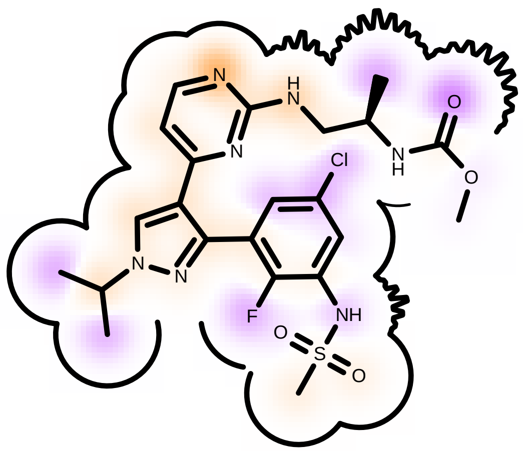

Score compared to other molecules

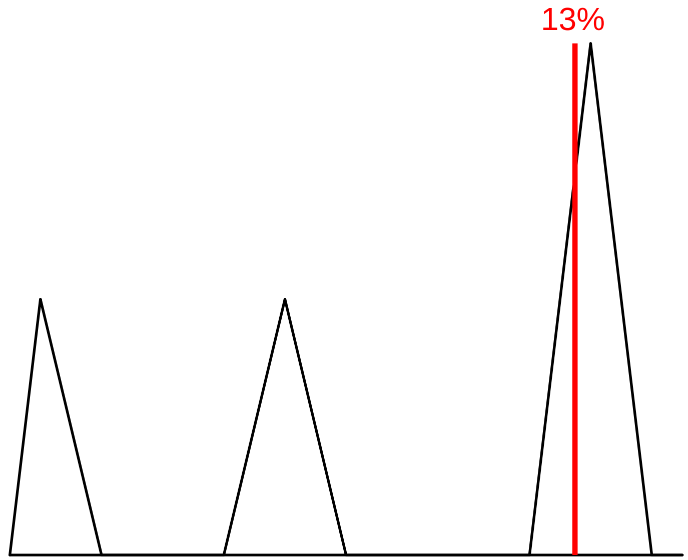

Better scores Worse scores

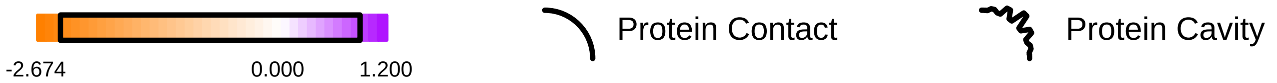

#### Residue Fingerprint

| | |
|---|---|
| ALA481A | ASN580A |
| ASP594A | CYS532A |
| GLN530A | GLU533A |
| GLY464A | GLY466A |
| GLY534A | GLY593A |
| ILE463A | ILE527A |
| LEU505A | LEU514A |
| LYS483A | PHE468A |
| PHE583A | PHE595A |
| SER465A | SER535A |
| SER536A | THR529A |
| TRP531A | VAL471A |

| | |
|---|---|
| Acceptor | Donor |
| Metal | Contact |

#### Shape -10.86

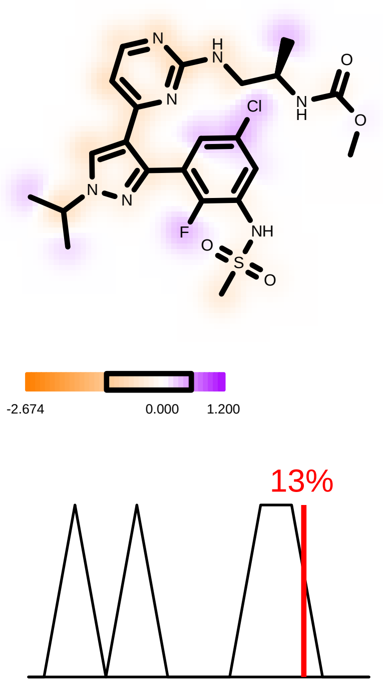

#### Hydrogen Bond -4.64

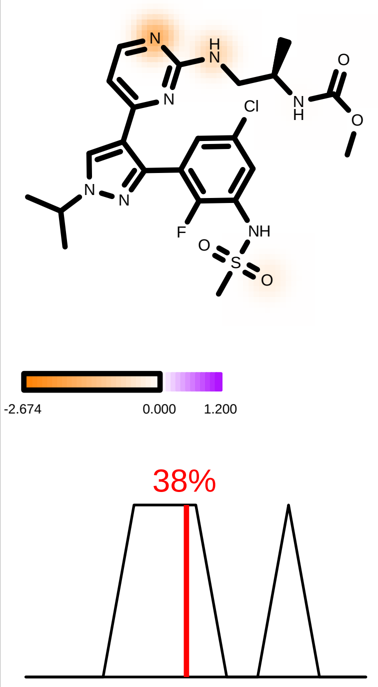

#### Protein Desolvation 3.16

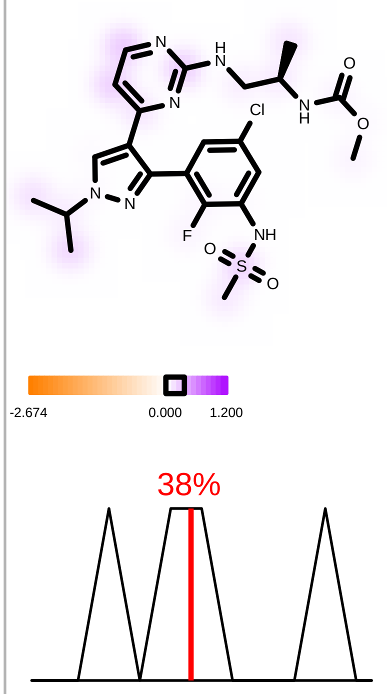

#### Ligand Desolvation 4.54

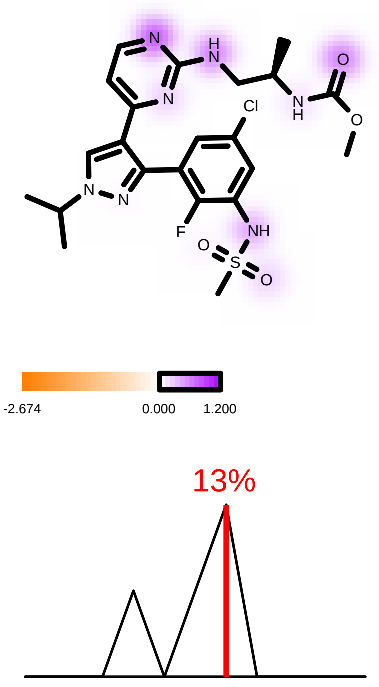

| | |
|---|---|
| Molecule Name | Cluster 14, 1 of 118 196 |
| Molecular Weight | 539.7 |
| XLogP | 4.9 |
| PSA | 110.9 |
| Heavy Atoms | 37 |
| Acceptor Count | 6 |
| Donor Count | 2 |
| Chelator Count | 3 |

**PDB ID: 4XV2**

*(Compound I)*

### Total Score -13.62

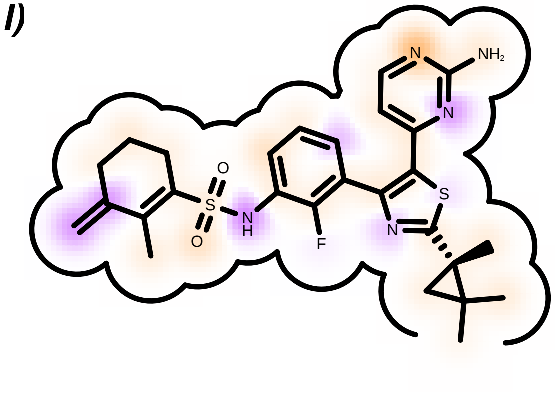

Score compared to other molecules

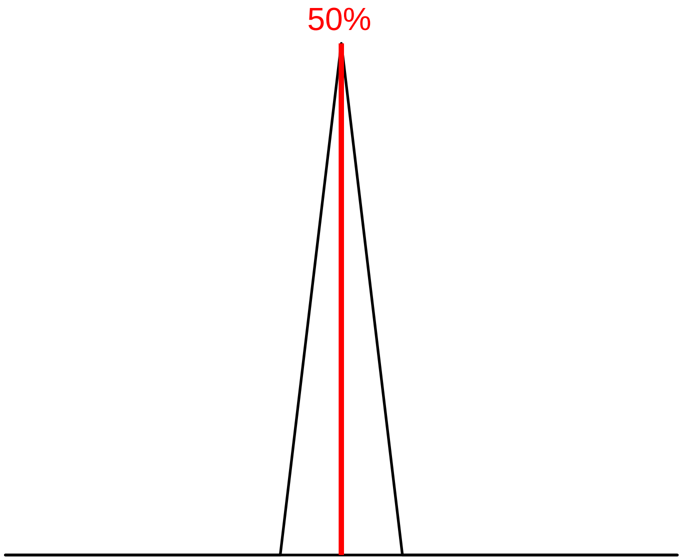

Better scores Worse scores

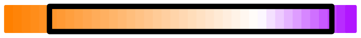

Protein Contact Protein Cavity

Residue Fingerprint

| | |
|---|---|
| ALA481A | ASN580A |
| **ASP594A** | **CYS532A** |
| GLN530A | GLY464A |
| GLY466A | ILE463A |
| LEU505A | LEU514A |
| **LYS483A** | PHE468A |
| PHE583A | PHE595A |
| SER465A | THR529A |
| TRP531A | VAL471A |
| VAL482A | |

| | |
|---|---|
| Acceptor | Donor |
| Metal | Contact |

#### Shape -18.99

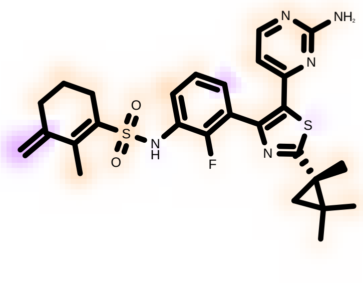

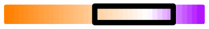

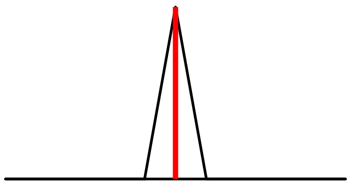

#### Hydrogen Bond -4.99

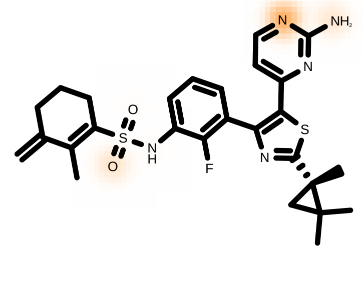

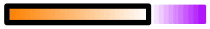

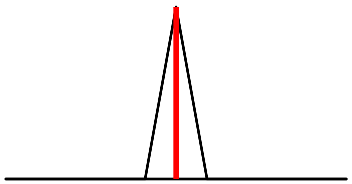

#### Protein Desolvation 4.85

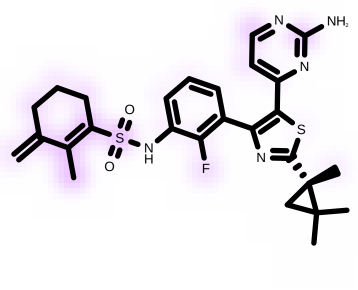

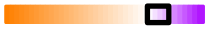

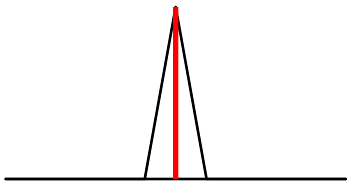

#### Ligand Desolvation 5.51

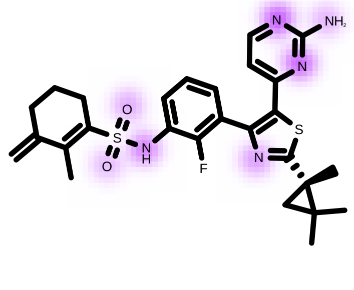

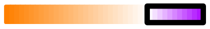

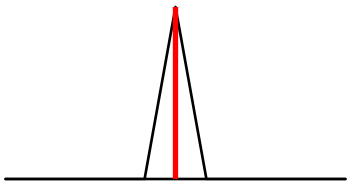

| | |
|---|---|
| Molecule Name | Molecule Name_122 |
| Molecular Weight | 509.7 |
| XLogP | 4.6 |
| PSA | 105.6 |
| Heavy Atoms | 35 |
| Acceptor Count | 5 |
| Donor Count | 1 |
| Chelator Count | 3 |

PDB ID: 4XV2

*(Compound II)*

### Total Score -14.33

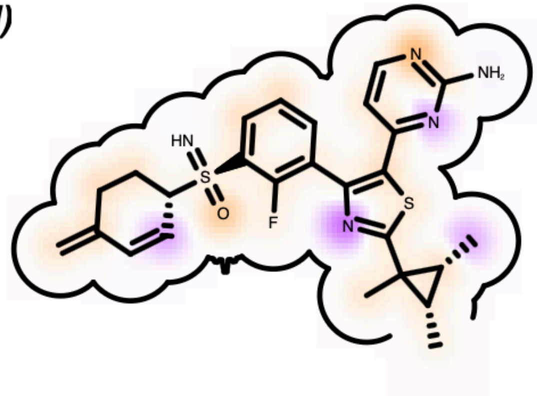

Score compared to other molecules

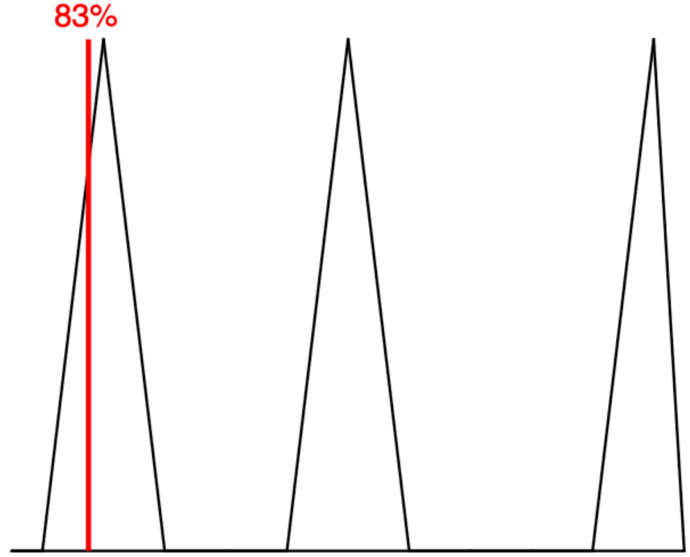

Better scores Worse scores

-2.416 0.000 1.200

Protein Contact

Protein Cavity

Residue Fingerprint

| | |
|---|---|
| ALA481A | ASP594A |
| CYS532A | GLN530A |
| GLY464A | GLY466A |
| GLY593A | ILE463A |
| ILE513A | ILE527A |
| LEU505A | LEU514A |
| LYS483A | PHE468A |
| PHE516A | PHE583A |
| PHE595A | SER465A |
| THR508A | THR529A |
| TRP531A | VAL471A |

Acceptor Donor
Metal Contact

#### Shape -18.12

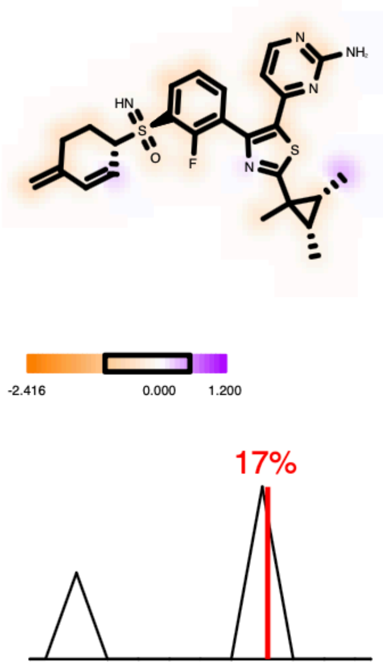

#### Hydrogen Bond -4.35

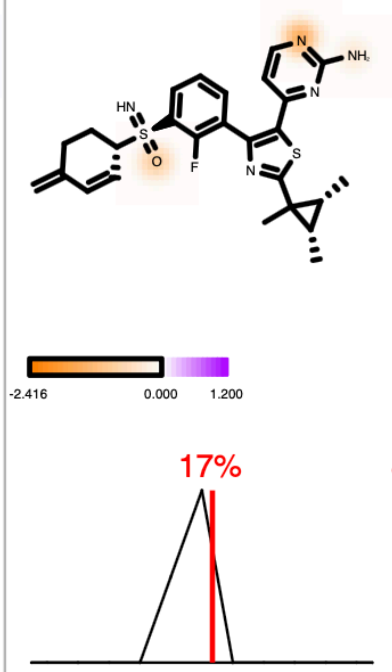

#### Protein Desolvation 3.85

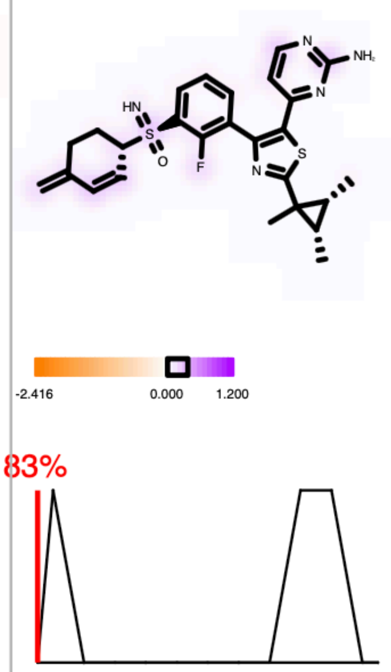

#### Ligand Desolvation 4.28

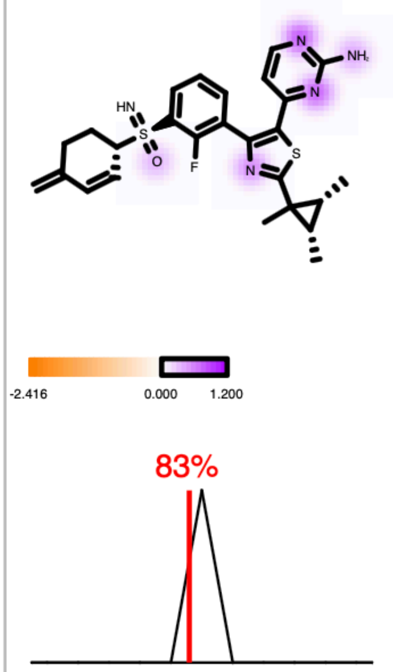

| | |
|---|---|
| Molecule Name | Vemurafenib_200 |
| Molecular Weight | 489.9 |
| XLogP | 5.4 |
| PSA | 91.9 |
| Heavy Atoms | 33 |
| Acceptor Count | 4 |
| Donor Count | 2 |
| Chelator Count | 1 |

PDB ID: 3OG7

### Total Score -17.67

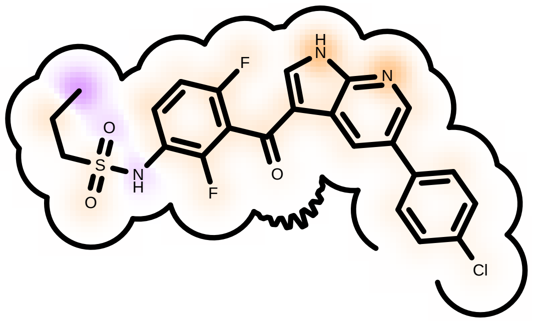

Score compared to other molecules

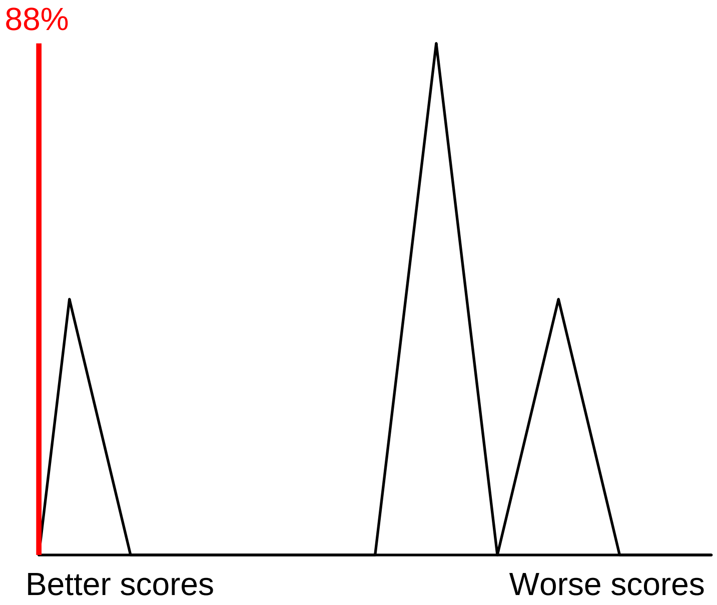

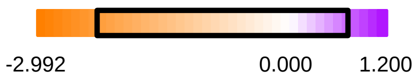

Protein Contact

Protein Cavity

Residue Fingerprint

| | |
|---|---|
| ALA481A | ASN580A |
| ASN581A | ASP594A |
| CYS532A | GLN530A |
| GLU501A | GLY464A |
| GLY593A | GLY596A |
| HIS539A | ILE463A |
| ILE527A | LEU505A |
| LEU514A | LEU515A |
| LYS483A | LYS578A |
| PHE468A | PHE583A |
| PHE595A | SER465A |
| SER535A | SER536A |
| THR529A | TYR538A |
| VAL471A | |

Acceptor Donor
Metal Contact

#### Shape -20.71

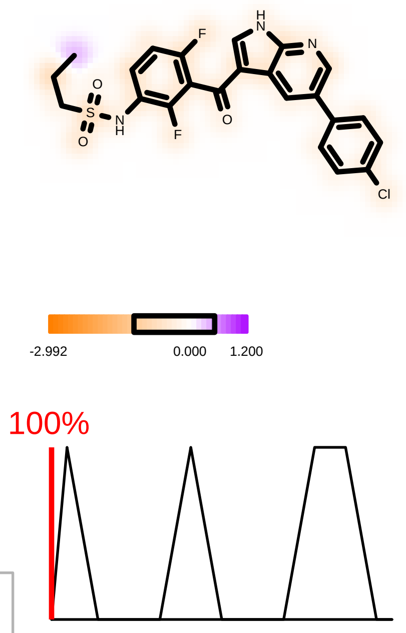

#### Hydrogen Bond -6.28

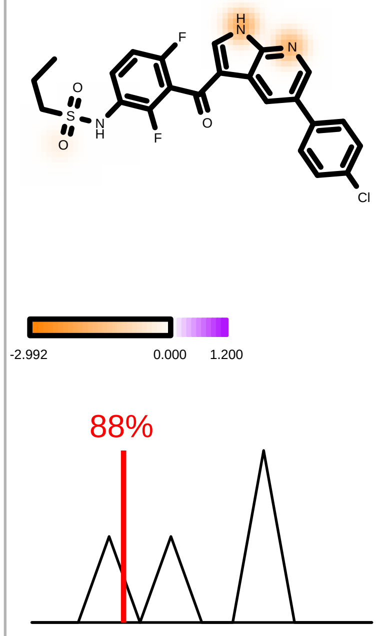

#### Protein Desolvation 4.70

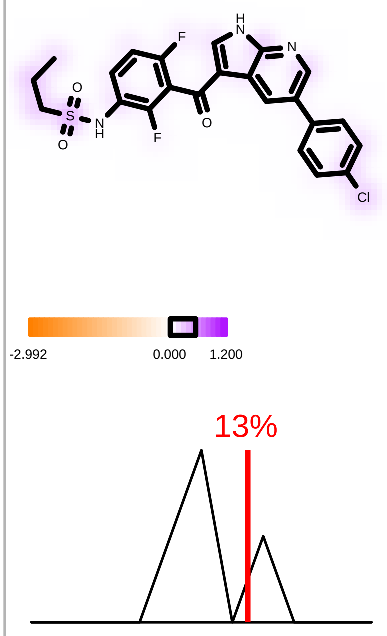

#### Ligand Desolvation 4.62

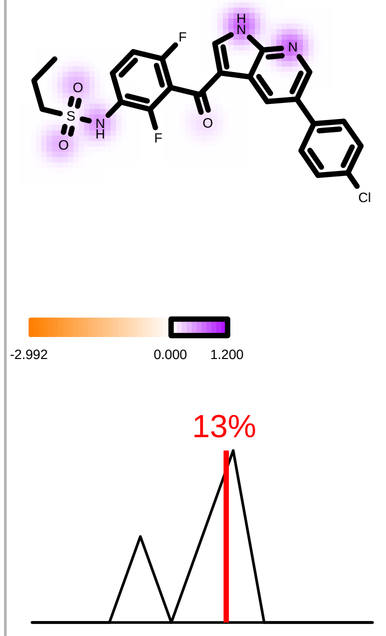

| | |
|---|---|
| Molecule Name | Molecule Name 139 |
| Molecular Weight | 519.6 |
| XLogP | 5.0 |
| PSA | 110.9 |
| Heavy Atoms | 35 |
| Acceptor Count | 6 |
| Donor Count | 2 |
| Chelator Count | 3 |

*(Dabrafenib)*

PDB ID: 3OG7

### Total Score -11.27

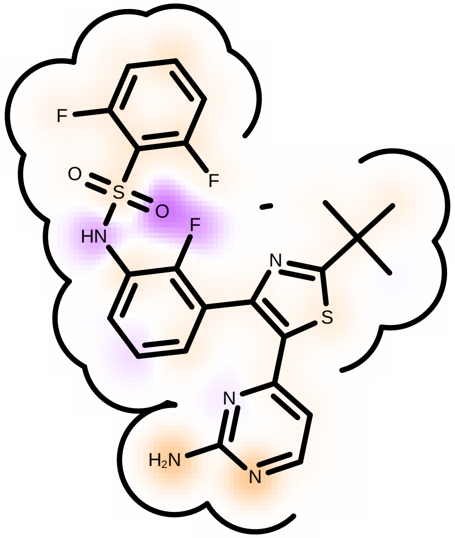

Score compared to other molecules

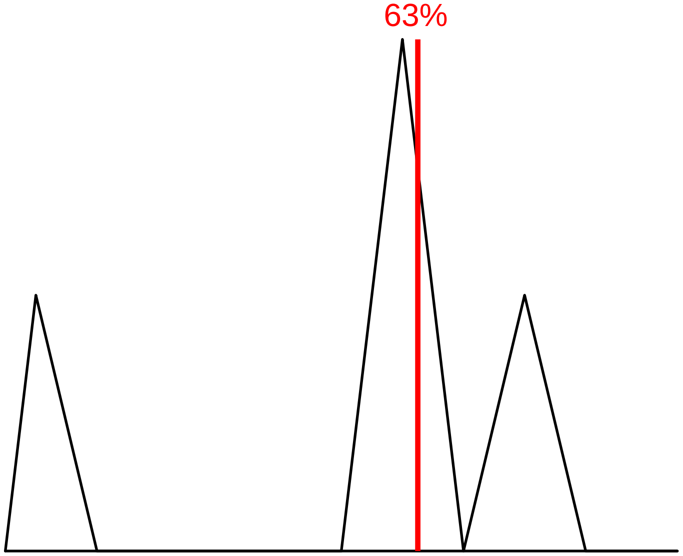

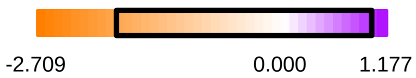

Protein Contact

Protein Cavity

#### Residue Fingerprint

| | |
|---|---|
| ALA481A | ASN580A |
| ASN581A | ASP594A |
| CYS532A | GLN530A |
| GLU501A | GLY464A |
| GLY593A | GLY596A |
| HIS539A | ILE463A |
| ILE527A | LEU505A |
| LEU514A | LEU515A |
| LYS483A | LYS578A |
| PHE468A | PHE583A |
| PHE595A | SER465A |
| SER535A | SER536A |
| THR529A | TYR538A |
| VAL471A | |

| | |
|---|---|
| Acceptor | Donor |
| Metal | Contact |

Shape -13.20

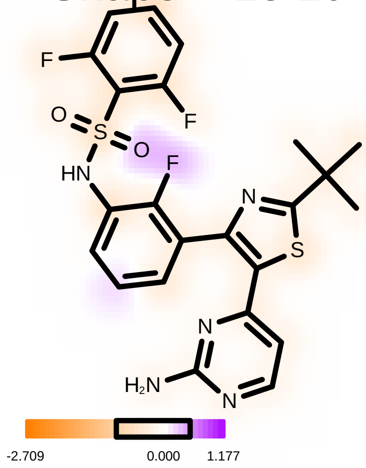

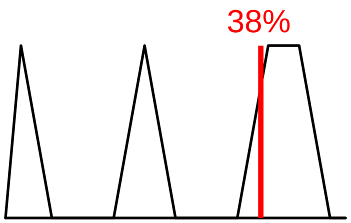

Hydrogen Bond -5.23

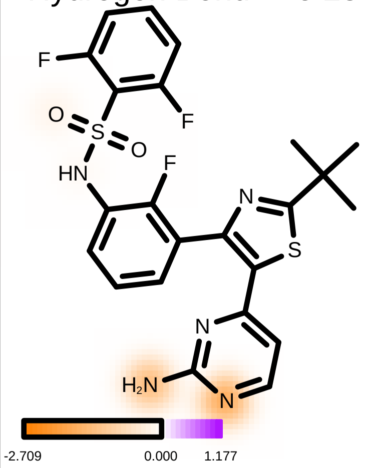

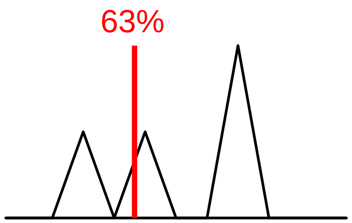

Protein Desolvation 3.07

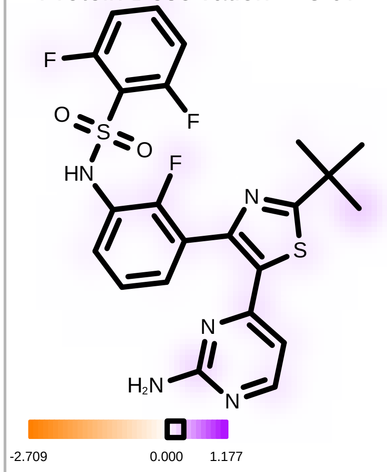

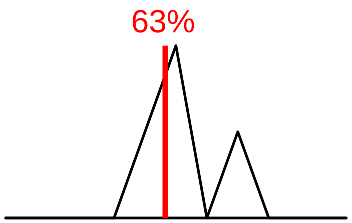

Ligand Desolvation 4.09

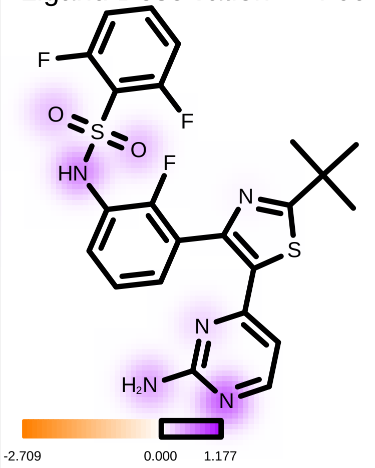

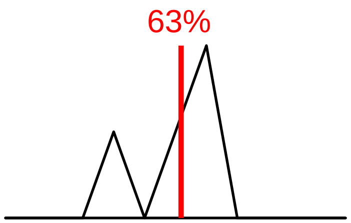

| | |
|---|---|
| Molecule Name | Sorafenib_165 |
| Molecular Weight | 464.8 |
| XLogP | 3.7 |
| PSA | 92.4 |
| Heavy Atoms | 32 |
| Acceptor Count | 4 |
| Donor Count | 3 |
| Chelator Count | 1 |

PDB ID: 3OG7

### Total Score -11.18

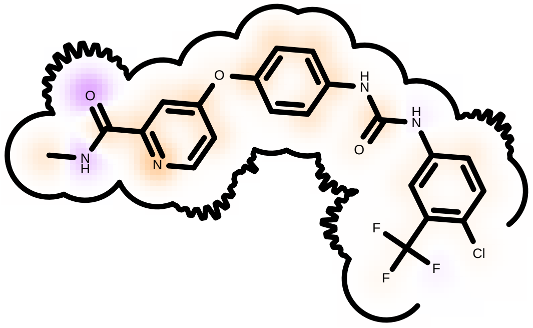

Score compared to other molecules

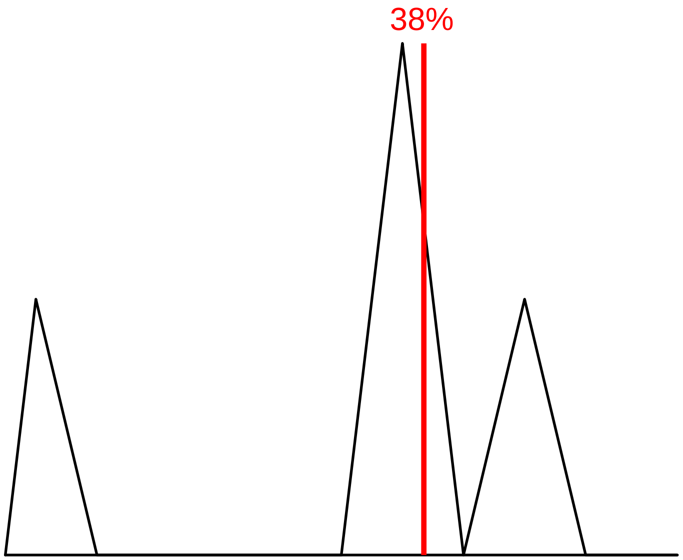

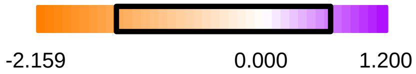

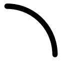 Protein Contact

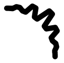 Protein Cavity

Residue Fingerprint

| | |
|---|---|
| ALA481A | ASN580A |
| ASN581A | ASP594A |
| CYS532A | GLN530A |
| GLU501A | GLY464A |
| GLY593A | **GLY596A** |
| HIS539A | ILE463A |
| ILE527A | LEU505A |
| LEU514A | LEU515A |
| LYS483A | LYS578A |
| PHE468A | PHE583A |
| PHE595A | SER465A |
| SER535A | SER536A |
| THR529A | TYR538A |
| VAL471A | |

| | |
|---|---|
| Acceptor | Donor |
| Metal | Contact |

#### Shape -16.71

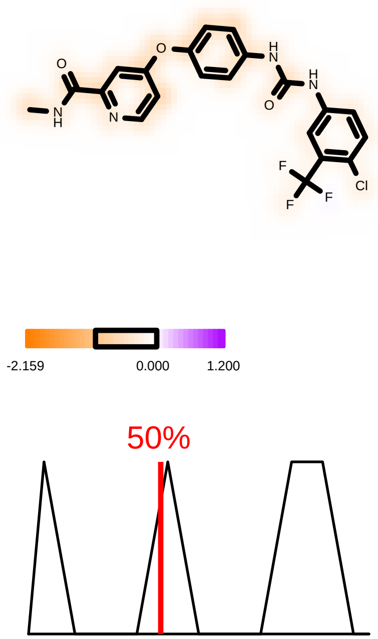

#### Hydrogen Bond -2.17

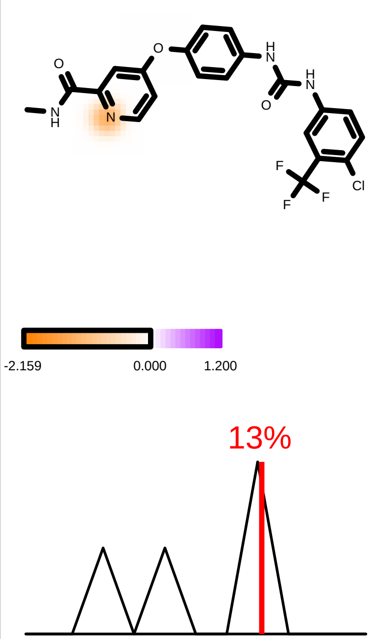

#### Protein Desolvation 3.14

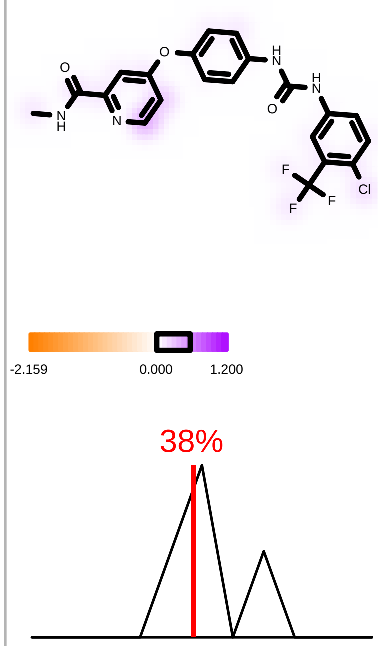

#### Ligand Desolvation 4.56

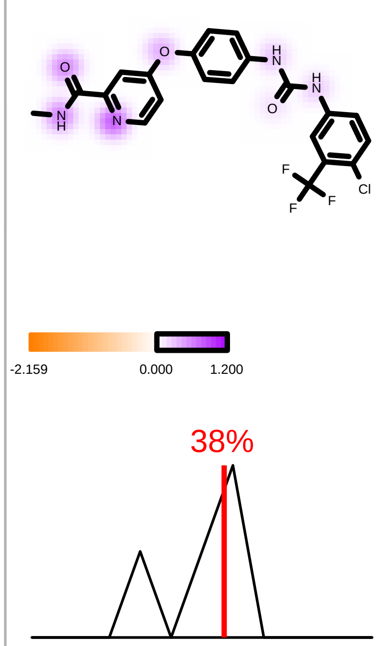

| | |
|---|---|
| Molecule Name | Molecule Name 43 |
| Molecular Weight | 540.0 |
| XLogP | 2.6 |
| PSA | 140.1 |
| Heavy Atoms | 36 |
| Acceptor Count | 6 |
| Donor Count | 3 |
| Chelator Count | 3 |

PDB ID: 3OG7

*(Encorafenib)*

### Total Score -9.36

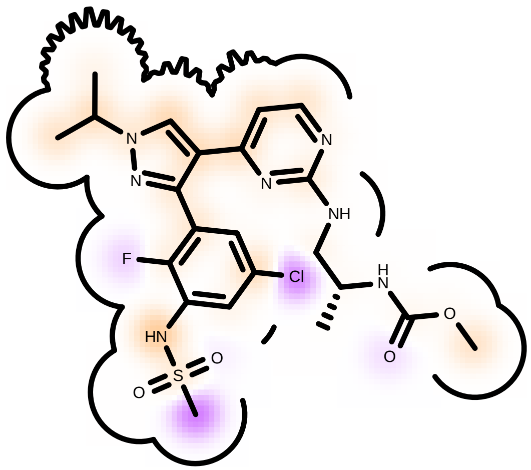

Score compared to other molecules

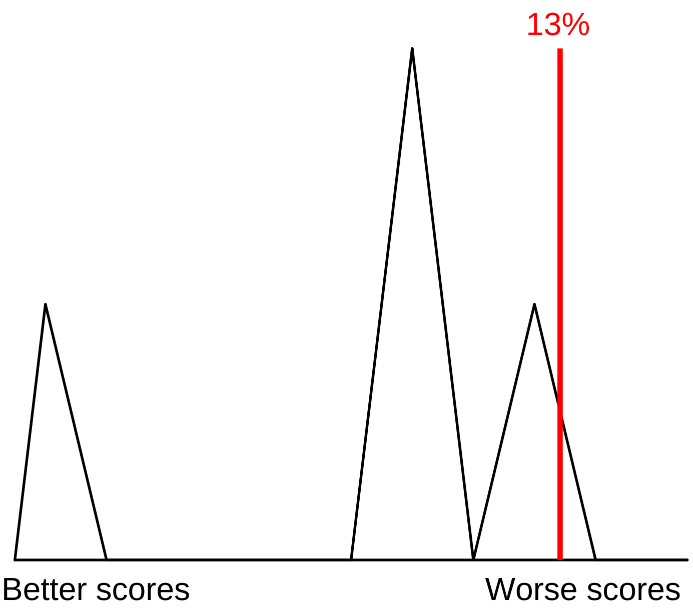

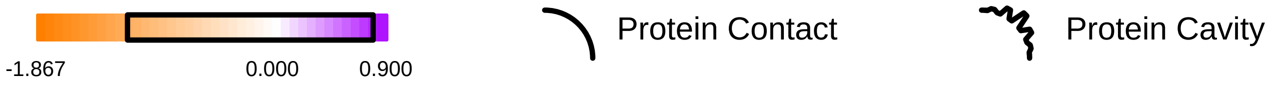

Residue Fingerprint

| | |
|---|---|
| ALA481A | ASN580A |
| ASN581A | **ASP594A** |
| CYS532A | GLN530A |
| GLU501A | GLY464A |
| GLY593A | GLY596A |
| HIS539A | ILE463A |
| ILE527A | LEU505A |
| LEU514A | LEU515A |
| LYS483A | LYS578A |
| PHE468A | PHE583A |
| PHE595A | SER465A |
| SER535A | SER536A |
| THR529A | TYR538A |
| VAL471A | |

| | |
|---|---|
| Acceptor | Donor |
| Metal | Contact |

Shape -11.91

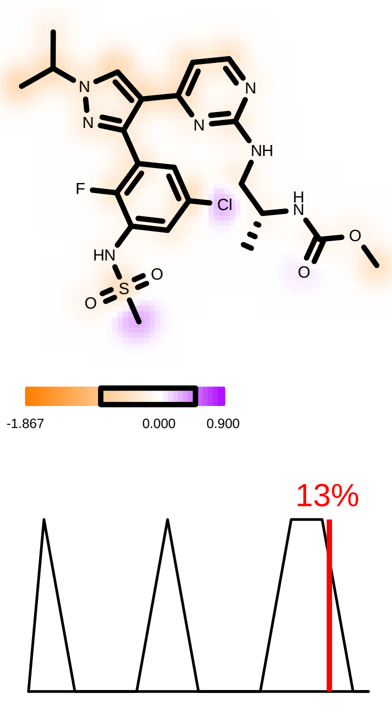

Hydrogen Bond -2.52

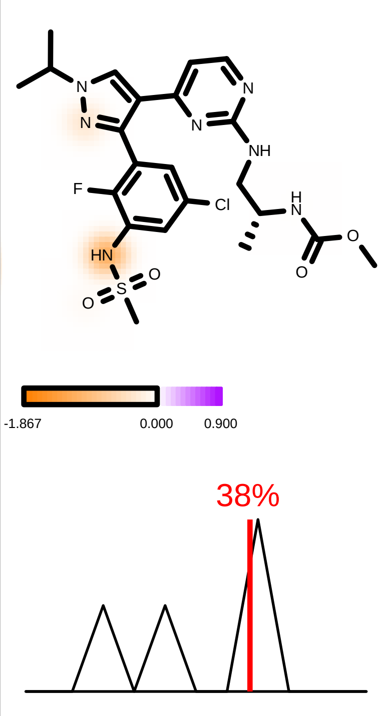

Protein Desolvation 2.60

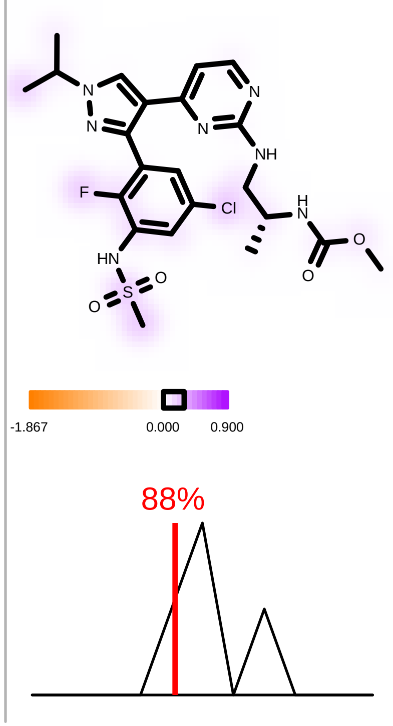

Ligand Desolvation 2.46

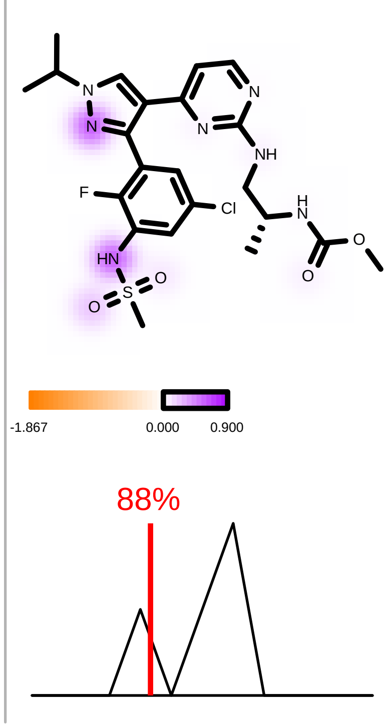

| | |
|---|---|
| Molecule Name | Cluster 14, 1 of 118 196 |
| Molecular Weight | 539.7 |
| XLogP | 4.9 |
| PSA | 110.9 |
| Heavy Atoms | 37 |
| Acceptor Count | 6 |
| Donor Count | 2 |
| Chelator Count | 3 |

PDB ID: 3OG7

### Total Score -12.88

*(Compound I)*

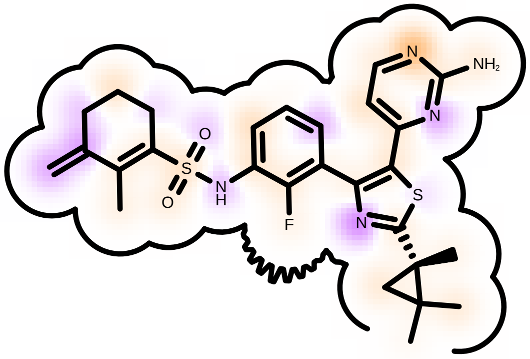

Score compared to other molecules

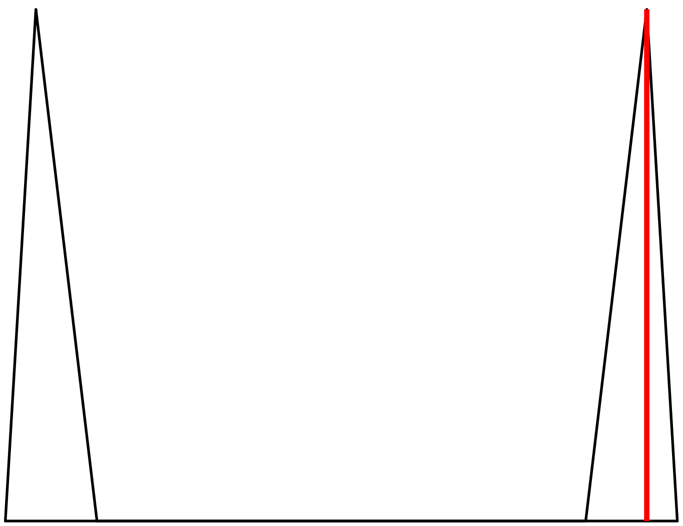

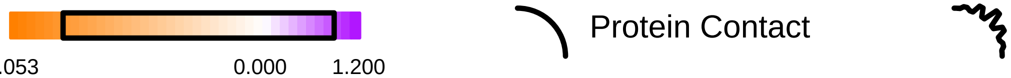

Residue Fingerprint

| | |
|---|---|
| ALA481A | ASN580A |
| ASN581A | ASP594A |
| CYS532A | GLN530A |
| GLY464A | GLY466A |
| GLY596A | ILE463A |
| ILE527A | LEU505A |
| LEU514A | LYS483A |
| PHE468A | PHE516A |
| PHE583A | PHE595A |
| SER465A | THR529A |
| TRP531A | VAL471A |
| VAL482A | |

Acceptor Donor
Metal Contact

#### Shape -17.78

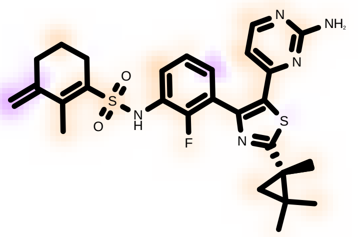

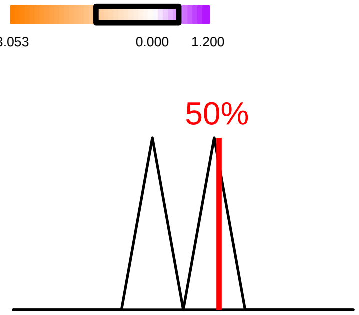

#### Hydrogen Bond -5.10

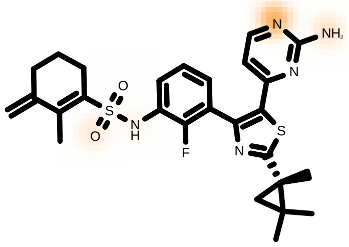

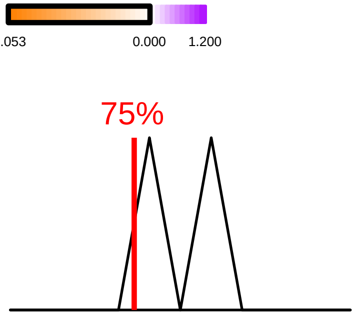

#### Protein Desolvation 4.42

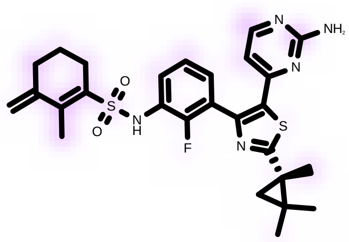

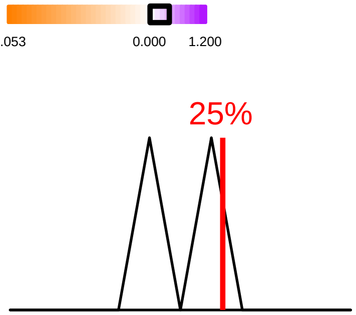

#### Ligand Desolvation 5.58

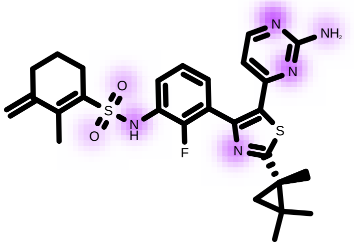

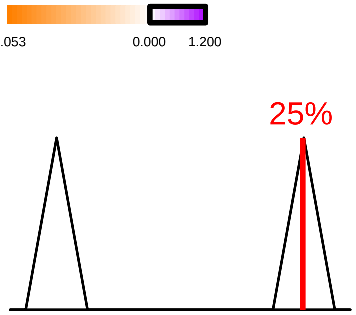

| | |
|---|---|
| Molecule Name | Molecule Name 122 |
| Molecular Weight | 509.7 |
| XLogP | 4.6 |
| PSA | 105.6 |
| Heavy Atoms | 35 |
| Acceptor Count | 5 |
| Donor Count | 1 |
| Chelator Count | 3 |

PDB ID: 3OG7

### Total Score -14.99

***(Compound II)***

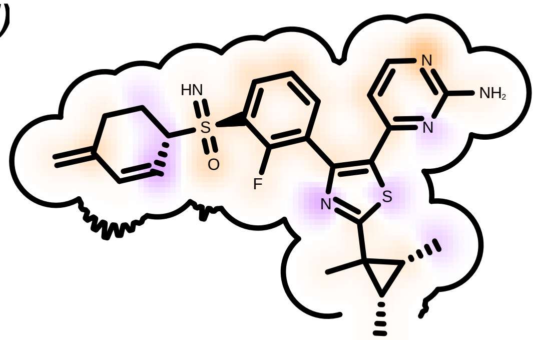

Score compared to other molecules

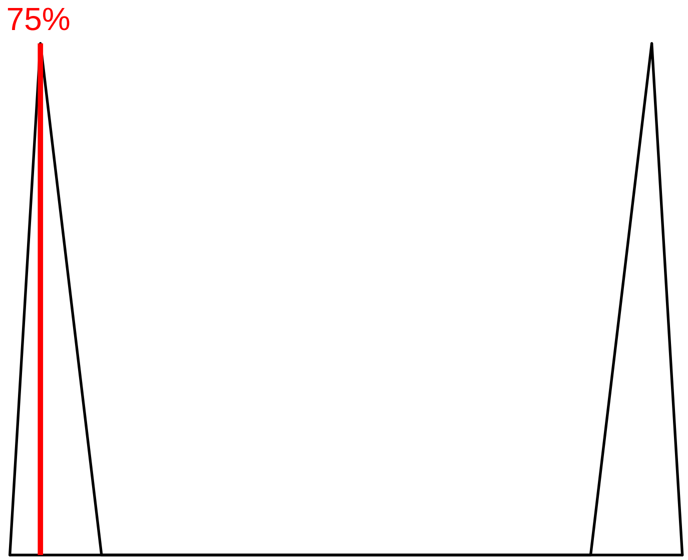

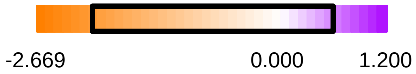

Protein Contact

Protein Cavity

Residue Fingerprint

| | |
|---|---|
| ALA481A | ASN580A |
| ASN581A | ASP594A |
| CYS532A | GLN530A |
| GLY464A | GLY466A |
| GLY596A | ILE463A |
| ILE527A | LEU505A |
| LEU514A | LYS483A |
| PHE468A | PHE516A |
| PHE583A | PHE595A |
| SER465A | THR529A |
| TRP531A | VAL471A |
| VAL482A | |

| | |
|---|---|
| Acceptor | Donor |
| Metal | Contact |

Shape -18.27

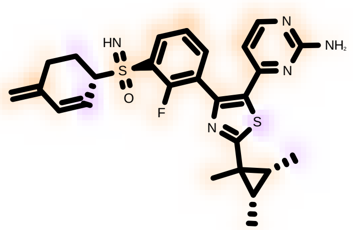

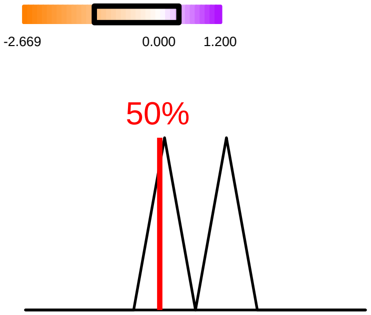

Hydrogen Bond -4.47

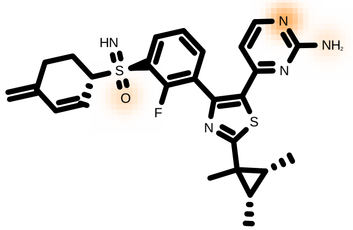

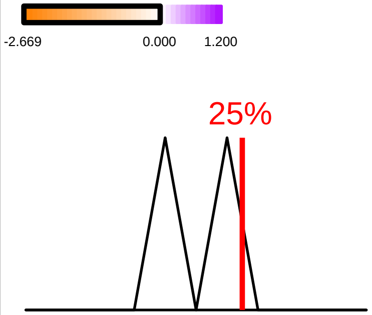

Protein Desolvation 3.85

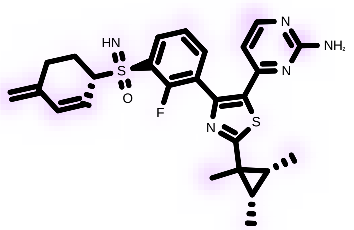

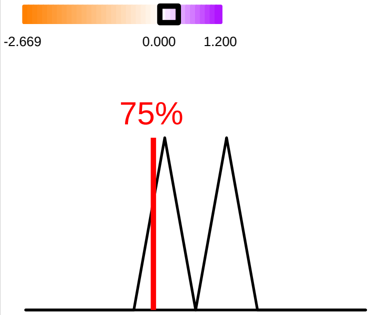

Ligand Desolvation 3.91

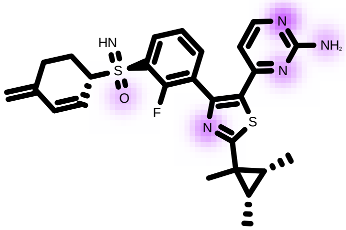

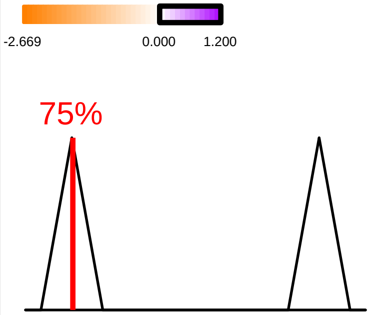

| | |
|---|---|
| Molecule Name | Vemurafenib_130 |
| Molecular Weight | 489.9 |
| XLogP | 5.4 |
| PSA | 91.9 |
| Heavy Atoms | 33 |
| Acceptor Count | 4 |
| Donor Count | 2 |
| Chelator Count | 1 |

PDB ID: 1UWH

### Total Score -18.30

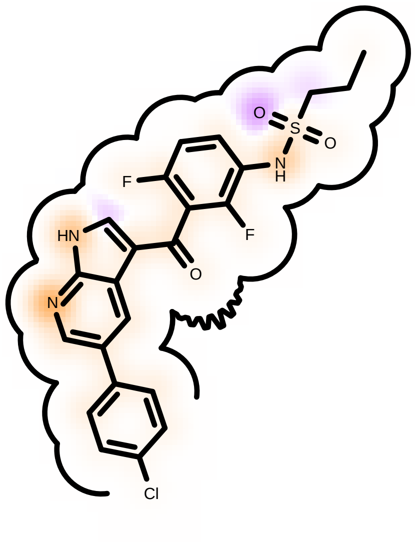

Score compared to other molecules

88%

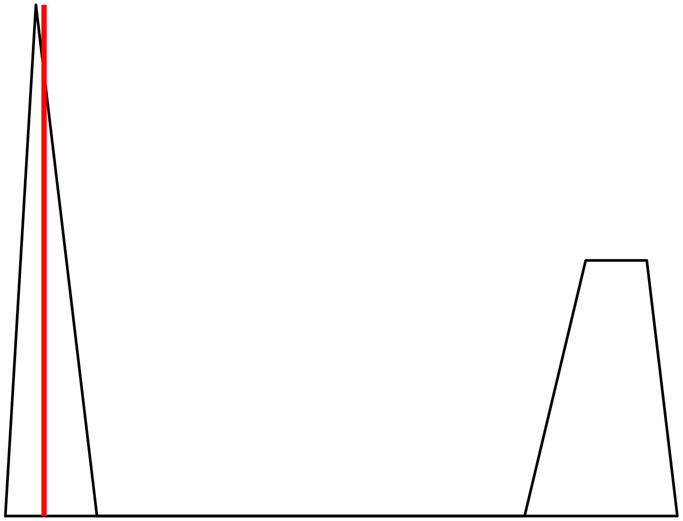

Better scores

Worse scores

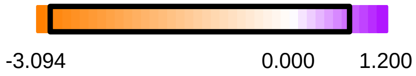

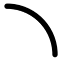 Protein Contact

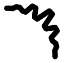 Protein Cavity

#### Residue Fingerprint

| | |
|---|---|
| ALA480B | ASN579B |
| ASP593B | CYS531B |
| GLN529B | GLU500B |
| GLY463B | GLY592B |
| HIS573B | ILE462B |
| ILE512B | ILE526B |
| ILE571B | ILE572B |
| LEU504B | LEU513B |
| LEU566B | LEU596B |
| LYS482B | NME599B |
| PHE582B | PHE594B |
| SER535B | THR507B |
| THR528B | TRP530B |
| VAL470B | VAL503B |

| | |
|---|---|
| Acceptor | Donor |
| Metal | Contact |

#### Shape -18.94

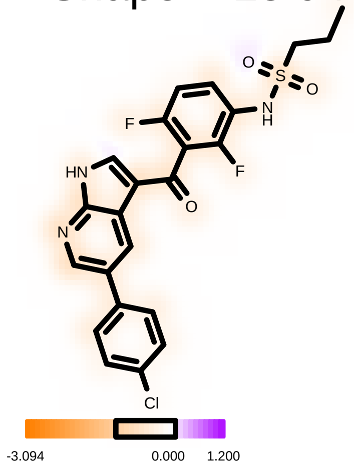

63%

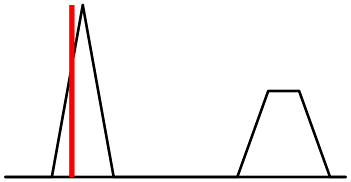

#### Hydrogen Bond -8.16

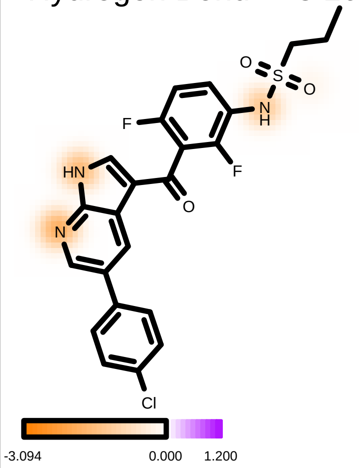

88%

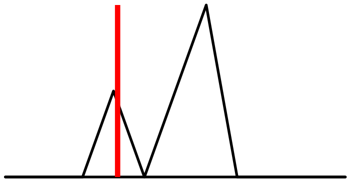

#### Protein Desolvation 4.26

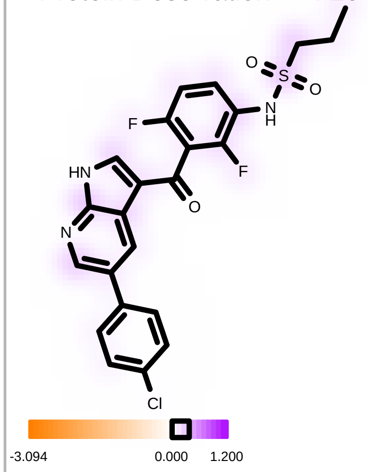

38%

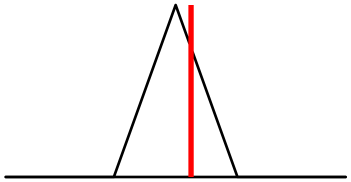

#### Ligand Desolvation 4.53

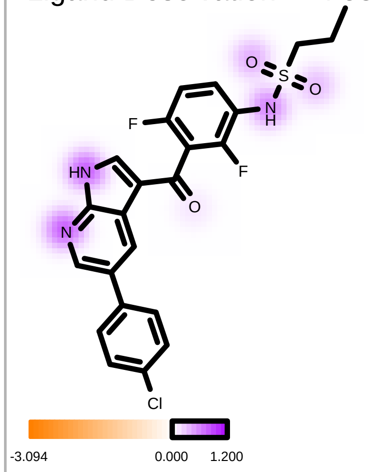

38%

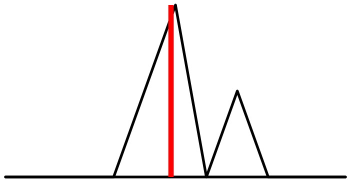

| | |
|---|---|
| Molecule Name | Sorafenib_13 |
| Molecular Weight | 464.8 |
| XLogP | 3.7 |
| PSA | 92.4 |
| Heavy Atoms | 32 |
| Acceptor Count | 4 |
| Donor Count | 3 |
| Chelator Count | 1 |

PDB ID: 1UWH

### Total Score -17.93

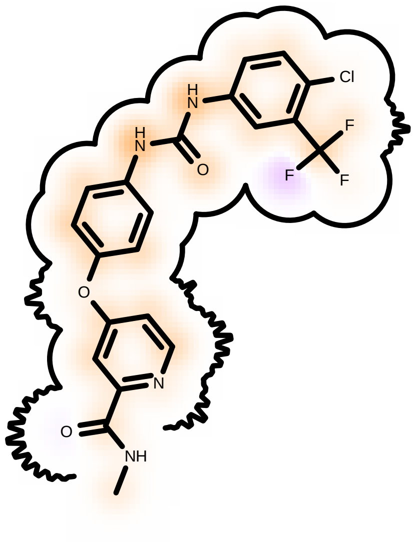

Score compared to other molecules

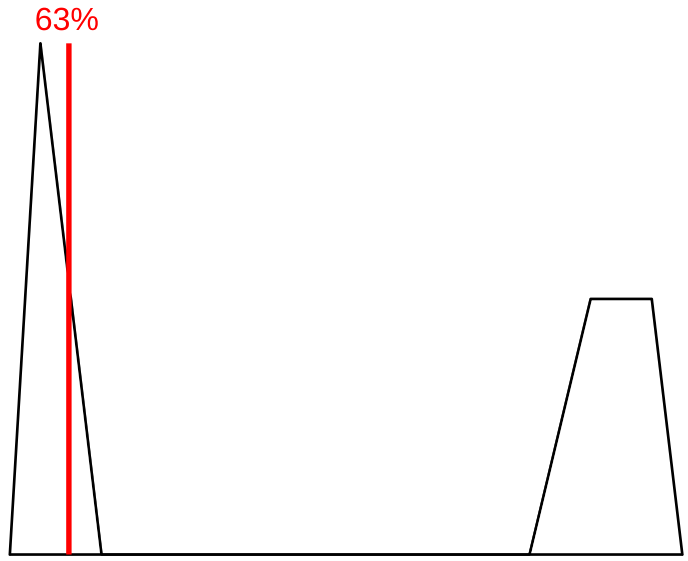

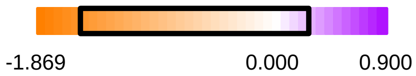

Protein Contact

Protein Cavity

Residue Fingerprint

| | |
|---|---|
| ALA480B | ASN579B |
| ASP593B | CYS531B |
| GLN529B | GLU500B |
| GLY463B | GLY592B |
| HIS573B | ILE462B |
| ILE512B | ILE526B |
| ILE571B | ILE572B |
| LEU504B | LEU513B |
| LEU566B | LEU596B |
| LYS482B | NME599B |
| PHE582B | PHE594B |
| SER535B | THR507B |
| THR528B | TRP530B |
| VAL470B | VAL503B |

| | |
|---|---|
| Acceptor | Donor |
| Metal | Contact |

Shape -19.07

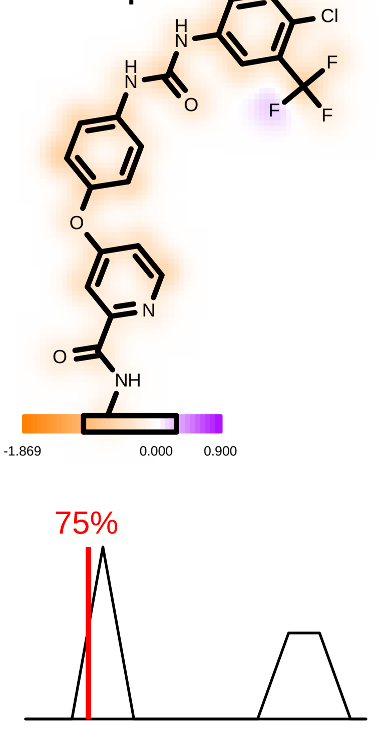

Hydrogen Bond -4.79

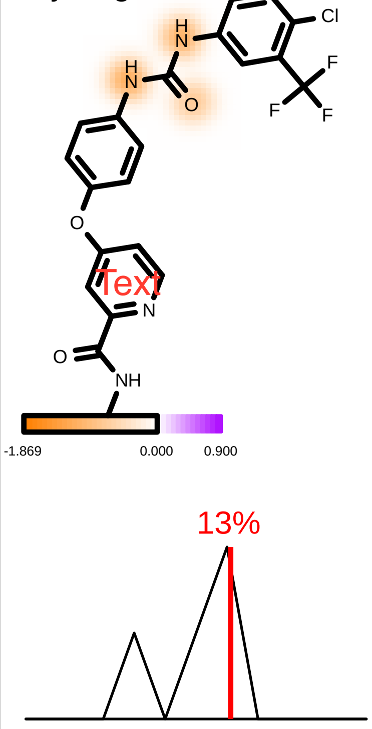

Protein Desolvation 2.48

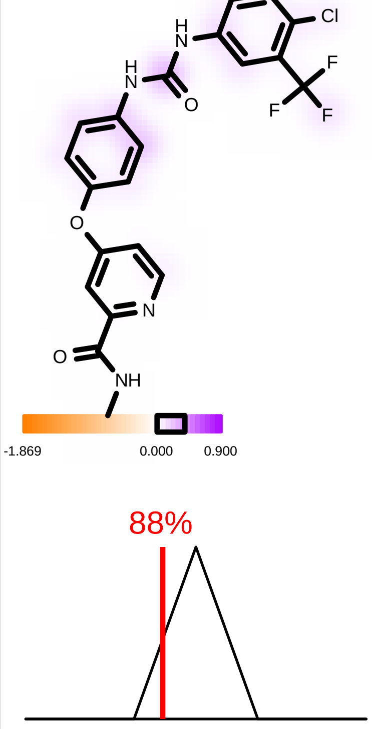

Ligand Desolvation 3.44

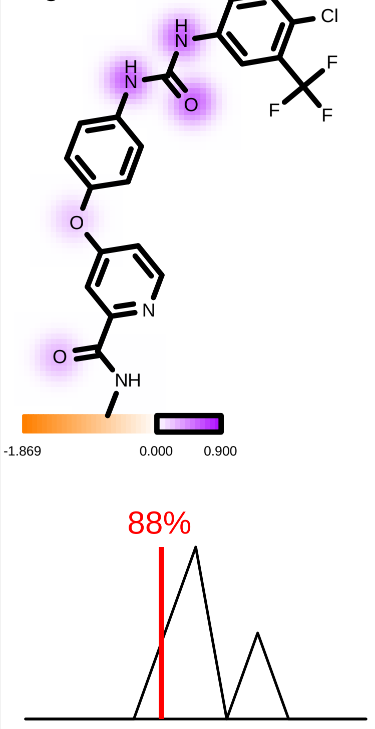

| | |
|---|---|
| Molecule Name | Molecule Name 85 |
| Molecular Weight | 519.6 |
| XLogP | 5.0 |
| PSA | 110.9 |
| Heavy Atoms | 35 |
| Acceptor Count | 6 |
| Donor Count | 2 |
| Chelator Count | 3 |

**(Dabrafenib)**

PDB ID: 1UWH

### Total Score -7.98

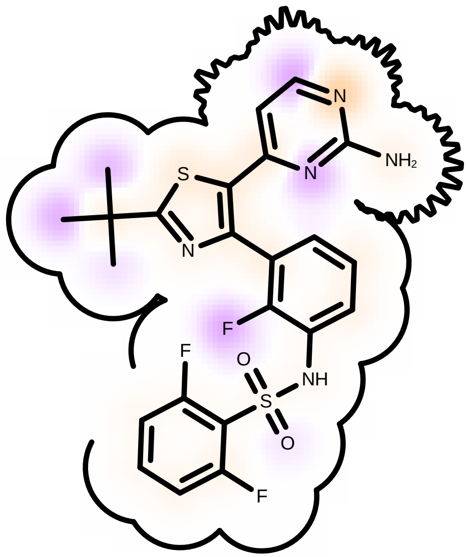

Score compared to other molecules

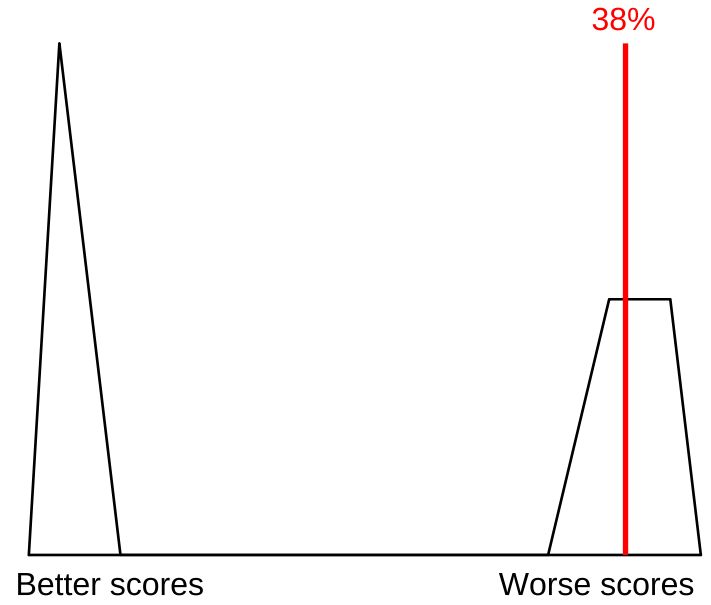

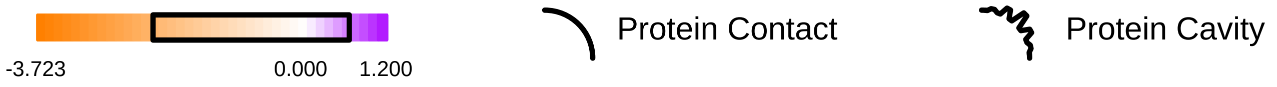

Residue Fingerprint

| | |
|---|---|
| ALA480B | ASN579B |
| ASP593B | CYS531B |
| GLN529B | **GLU500B** |
| GLY463B | GLY592B |
| HIS573B | ILE462B |
| ILE512B | ILE526B |
| ILE571B | ILE572B |
| LEU504B | LEU513B |
| LEU566B | LEU596B |
| **LYS482B** | NME599B |
| PHE582B | **PHE594B** |
| SER535B | THR507B |
| THR528B | TRP530B |
| VAL470B | VAL503B |

Acceptor Donor
Metal Contact

Shape -10.75

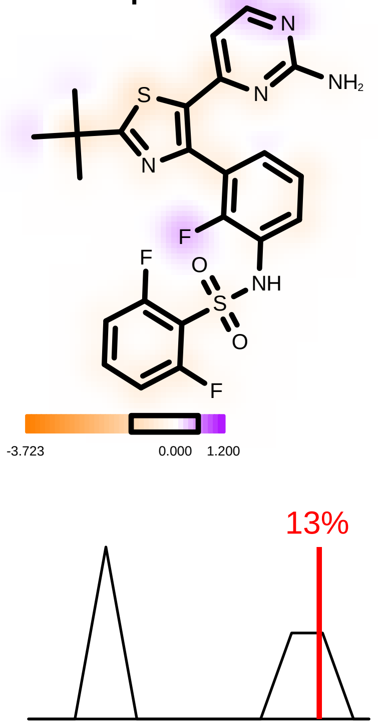

Hydrogen Bond -5.85

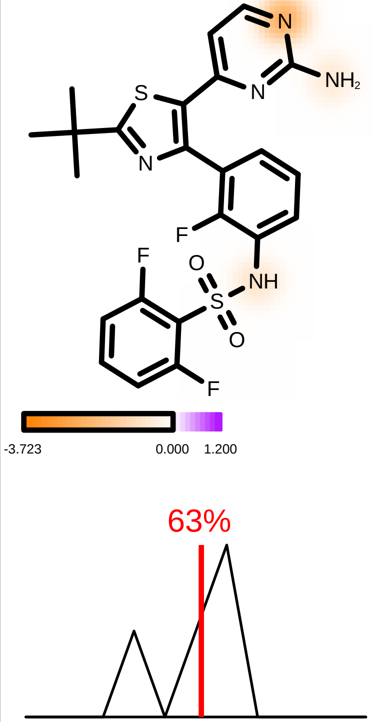

Protein Desolvation 4.32

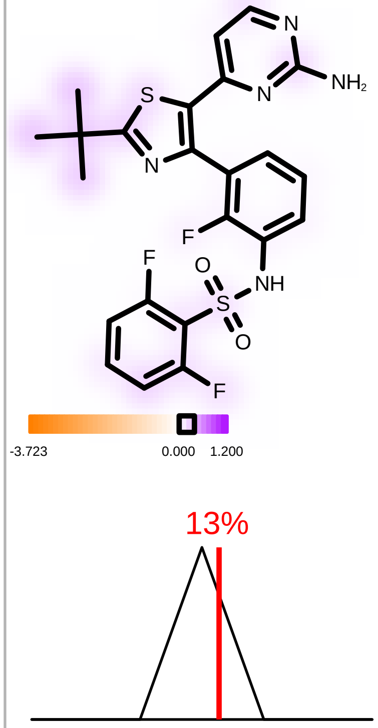

Ligand Desolvation 4.29

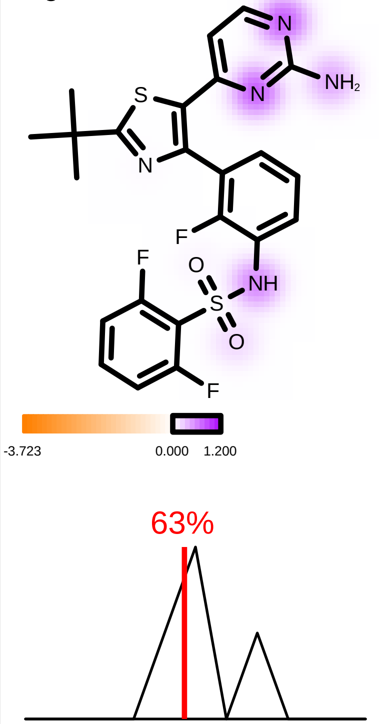

| | |
|---|---|
| Molecule Name | Molecule Name 88 |
| Molecular Weight | 540.0 |
| XLogP | 2.6 |
| PSA | 140.1 |
| Heavy Atoms | 36 |
| Acceptor Count | 6 |
| Donor Count | 3 |
| Chelator Count | 3 |

PDB ID: 1UWH

***(Encorafenib)***

### Total Score -7.00

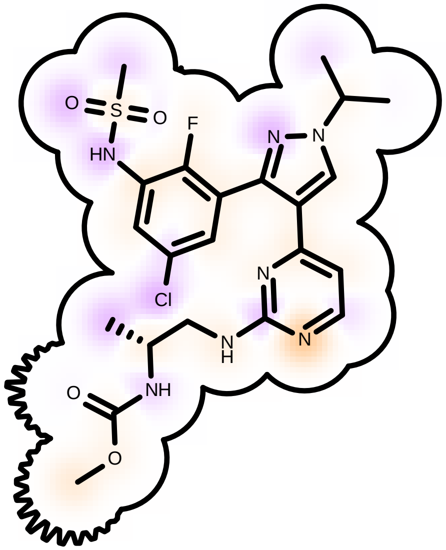

Score compared to other molecules

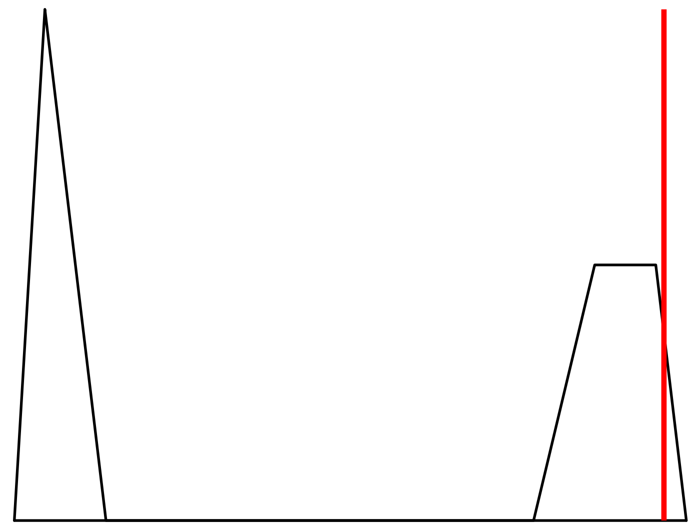

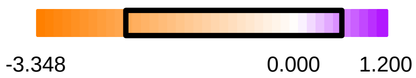

Protein Contact

Protein Cavity

Residue Fingerprint

| | |
|---|---|
| ALA480B | ASN579B |
| ASP593B | CYS531B |
| GLN529B | GLU500B |
| GLY463B | GLY592B |
| HIS573B | ILE462B |
| ILE512B | ILE526B |
| ILE571B | ILE572B |
| LEU504B | LEU513B |
| LEU566B | LEU596B |
| LYS482B | NME599B |
| PHE582B | PHE594B |
| SER535B | THR507B |
| THR528B | TRP530B |
| VAL470B | VAL503B |

| | |
|---|---|
| Acceptor | Donor |
| Metal | Contact |

Shape -11.83

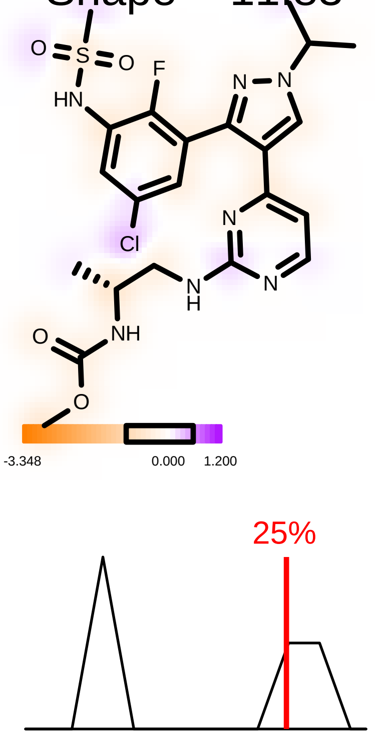

Hydrogen Bond -5.41

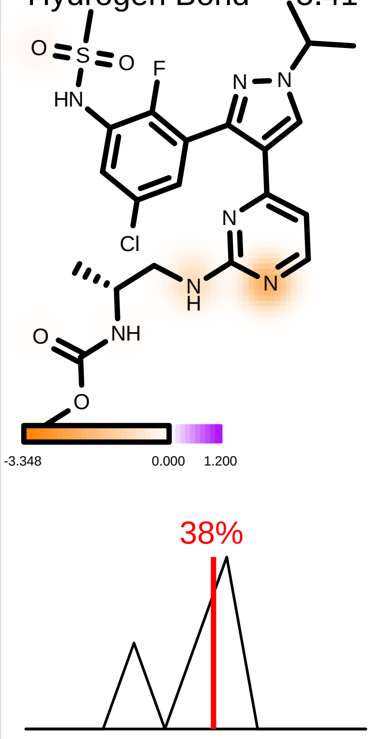

Protein Desolvation 3.72

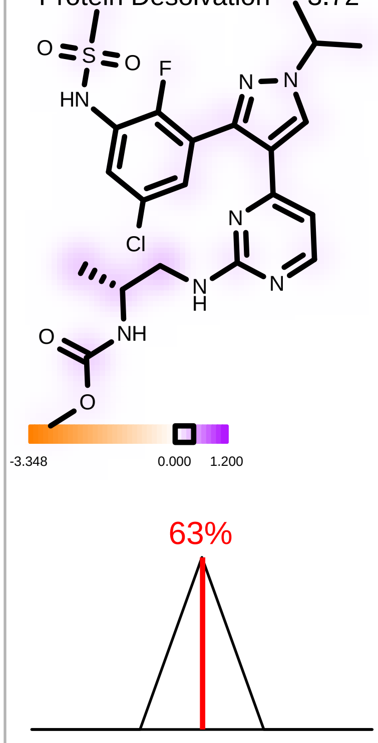

Ligand Desolvation 6.52

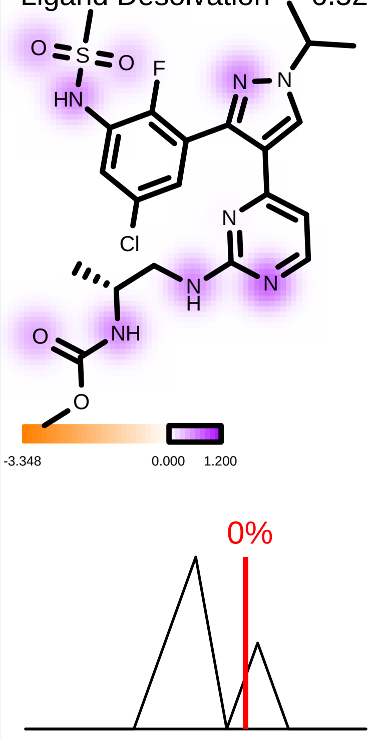

| | |
|---|---|
| Molecule Name | Cluster 14, 1 of 118 196 |
| Molecular Weight | 539.7 |
| XLogP | 4.9 |
| PSA | 110.9 |
| Heavy Atoms | 37 |
| Acceptor Count | 6 |
| Donor Count | 2 |
| Chelator Count | 3 |

PDB ID: 1UWH

*(Compound I)*

### Total Score -2.32

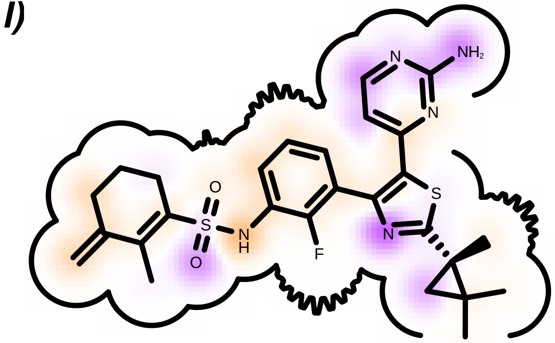

Score compared to other molecules

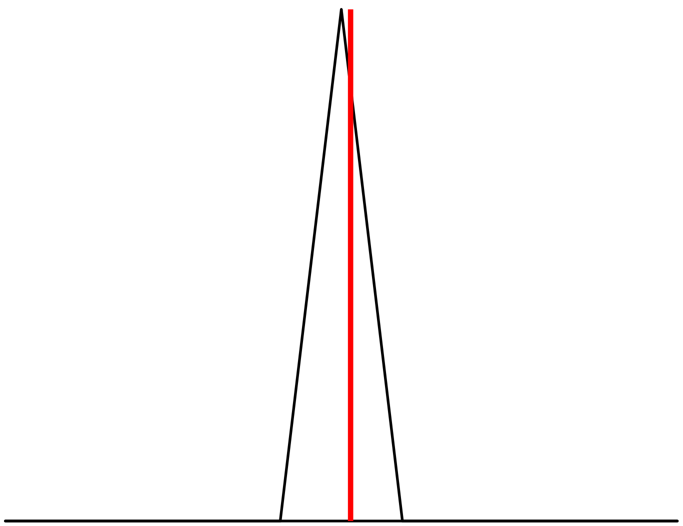

-1.726 0.000 1.377

Protein Contact

Protein Cavity

#### Residue Fingerprint

| | |
|---|---|
| ALA480B | ALA496B |
| ARG574B | ASN499B |
| ASN579B | ASP575B |
| **ASP593B** | CYS531B |
| GLN529B | **GLU500B** |
| GLY463B | GLY592B |
| GLY595B | HIS573B |
| ILE462B | ILE526B |
| LEU504B | LEU513B |
| LEU596B | LYS482B |
| NME599B | PHE582B |
| PHE594B | THR528B |
| THR598B | TRP530B |
| VAL470B | VAL501B |
| VAL503B | |

| | |
|---|---|
| Acceptor | Donor |
| Metal | Contact |

Shape -8.50

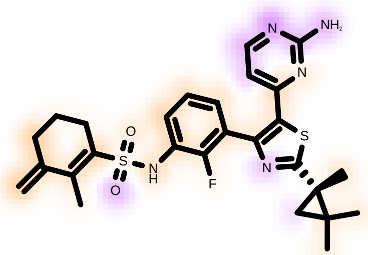

-1.726 0.000 1.377

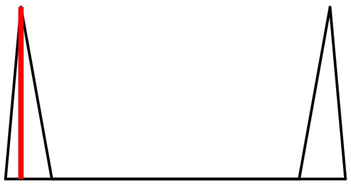

Hydrogen Bond -4.16

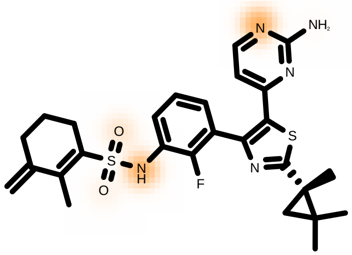

-1.726 0.000 1.377

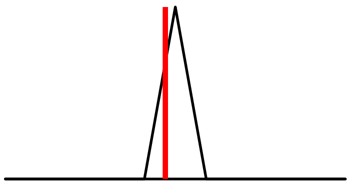

Protein Desolvation 5.68

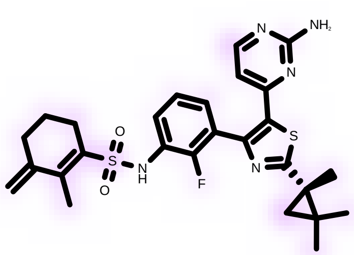

-1.726 0.000 1.377

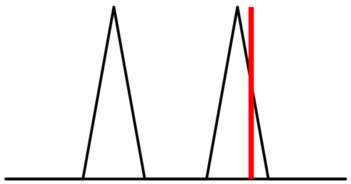

Ligand Desolvation 4.67

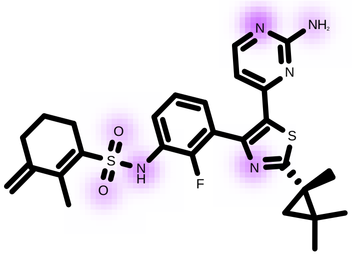

-1.726 0.000 1.377

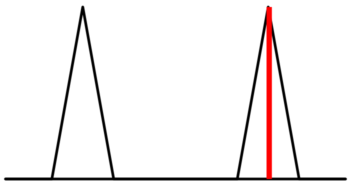

| | |
|---|---|
| Molecule Name | Molecule Name 122 |
| Molecular Weight | 509.7 |
| XLogP | 4.6 |
| PSA | 105.6 |
| Heavy Atoms | 35 |
| Acceptor Count | 5 |
| Donor Count | 1 |
| Chelator Count | 3 |

PDB ID: 1UWH

*(Compound II)*

### Total Score -2.39

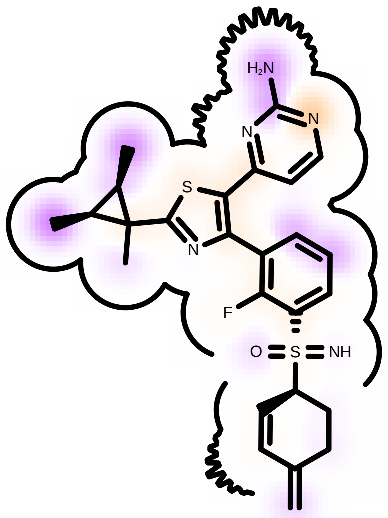

Score compared to other molecules

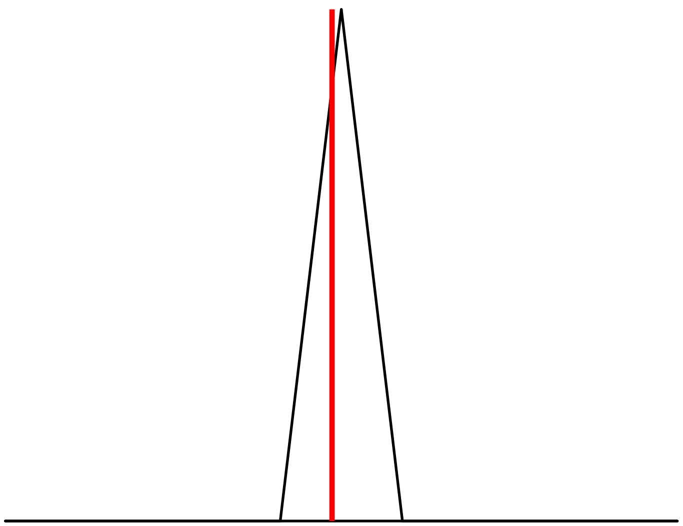

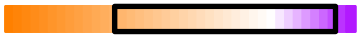

Protein Contact

Protein Cavity

#### Residue Fingerprint

| | |
|---|---|
| ALA480B | ALA496B |
| ARG574B | ASN499B |
| ASN579B | ASP575B |
| ASP593B | CYS531B |
| GLN529B | GLU500B |
| GLY463B | GLY592B |
| GLY595B | HIS573B |
| ILE462B | ILE526B |
| LEU504B | LEU513B |
| LEU596B | **LYS482B** |
| NME599B | PHE582B |
| PHE594B | THR528B |
| THR598B | TRP530B |
| VAL470B | VAL501B |
| VAL503B | |

| | |
|---|---|
| Acceptor | Donor |
| Metal | Contact |

Shape -6.10

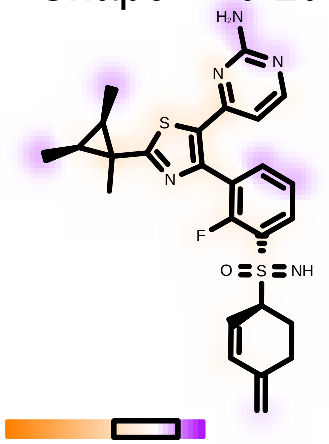

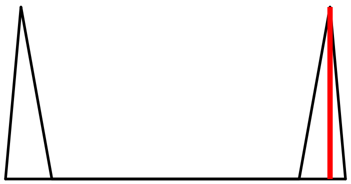

Hydrogen Bond -4.01

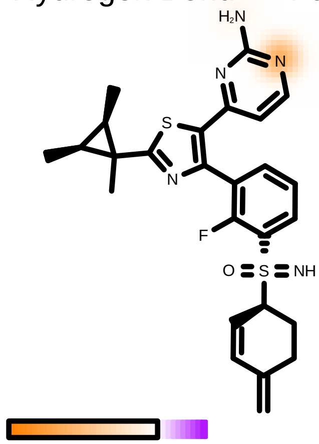

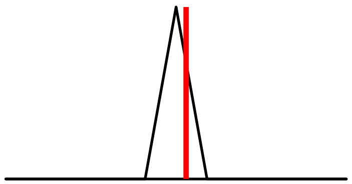

Protein Desolvation 4.51

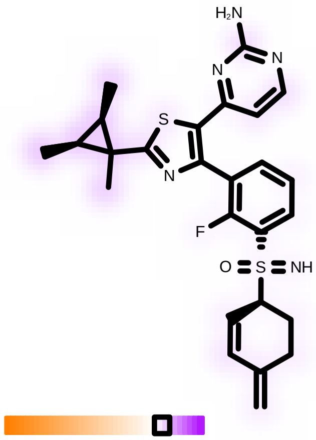

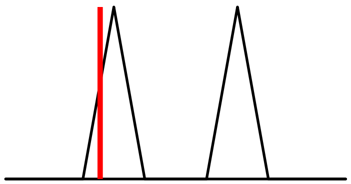

Ligand Desolvation 3.21

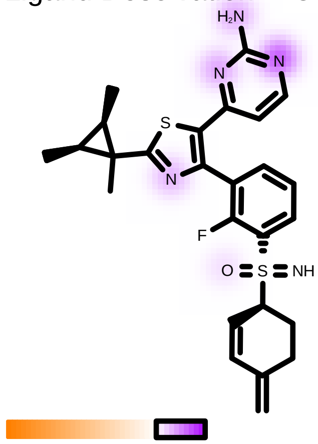

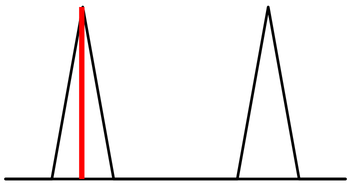
