## Supplementary material for "Computational Design of Two Novel BRAF V600E Inhibitors: Exploiting Sulfoximine Bioisosterism and Chiral Constraints to Evade Paradoxical Activation": File S2: BLAST Alignments

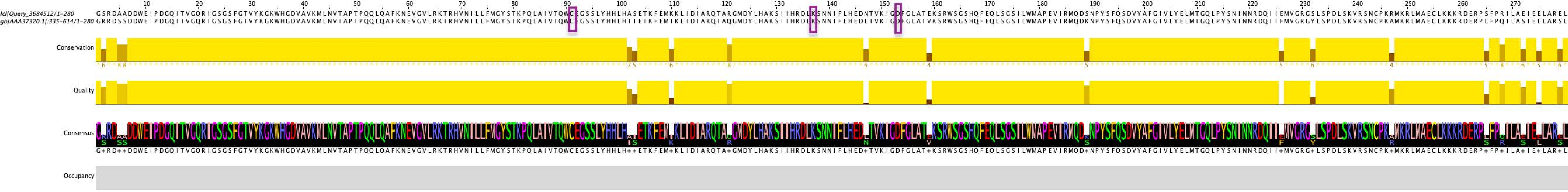
lcl|Query_3684512/1-280
gb|AAA37320.1|:335-614/1-280
Conservation
Quality
Consensus
Occupancy

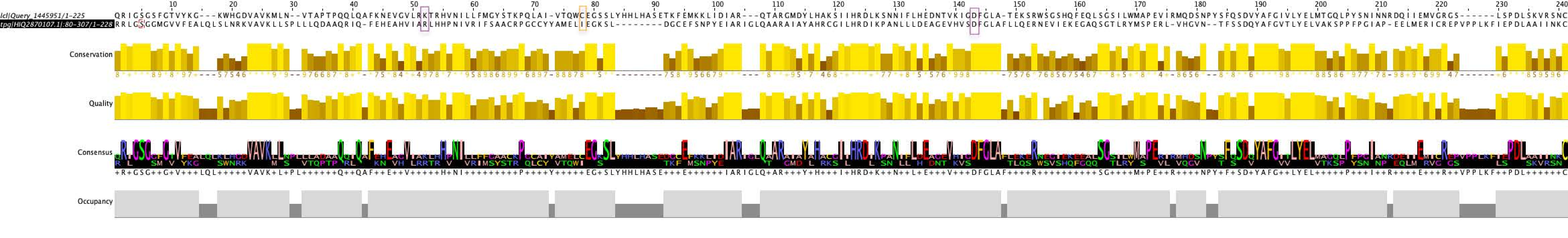

lcl|Query_1445951/1-225 QRIGSGSFGTVYKG---KWHGDVAVKMLN--VTAPTPQQLQAFKNEVGVLRKTRHVNILLFMGYSTKPQLAI-VTQWCEGSSLYHHLHASETKFEMKKLIDIAR---QTARGMDYLHAKSIIHRDLKSNNIFLHEDNTVKIGDFGLA-TEKSRWSGSHQFEQLSGSILWMAPEVIRMQDSNPYSFQSDVYAFGIVLYELMTGQLPYSNINNRDQIIEMVGRGS------LSPDLSKVRSNC
tpg|HIQ2870107.1|:80-307/1-228 RRLGSGGMGVVFEALQLSLNRKVAVKLLSPLLLQDAAQRIQ-FEHEAHVIARLHHPNIVRIFSAACRPGCCYYAMELIEGKSL--------DGCEFSNPYEIARIGLQAARAIAYAHRCGILHRDIKPANLLLDEAGEVHVSDFGLAFLLQERNEVIEKEGAQSGTLRYMSPERL-VHGVN--TFSSDQYAFGVTLYELVAKSPPFPGIAP-EELMERICREPVPPLKFIEPDLAAIINKC
Conservation
Quality
Consensus
Occupancy

NCBI Multiple Sequence Alignment Viewer, Version 1.26.0

| Sequence ID | | Start | Alignment | End | Organism |
|---|---|---|---|---|---|
| Query_6052307 | (+) | 96 | IVTQWCEGSSLYHHLHASETKFEM--KKLIDIARQTARGMDYLHAKSIIHRDLKSNNIFLHEDNTVKIGDFGLATEKSRWSGSHQFEQLSGSILWMAPEVIRMQDSNPYSFQSDVYAFGIVLYELMTGQLPY | 225 | |
| MCD0886487.1 | (+) | 32 | LVMEYVKGMDLKRYI---QTHFPIPYSTVVDITQQILSAVAMAHEHRIIHRDLKPQNILIDEHGTVKITDFGIAIALSETSIT-QTNTMLGSVHYLSPEQAR---GSMATNQSDIYAVGIILYEMLTGNVPF | 156 | Staphylococcus aureus |
