## Supplementary material for "Computational Design of Two Novel BRAF V600E Inhibitors: Exploiting Sulfoximine Bioisosterism and Chiral Constraints to Evade Paradoxical Activation": File S3: Methodological Transparency

### Supplementary Information: Methodological Transparency and AI-Assisted Copyediting

*"The current document serves as a proactive declaration of human agency, as well as a practical reference for the responsible application of AI tools in academic copyediting."*

**Parameters of AI Usage in the Current Study:**

***AI use was strictly confined to editorial support.*** All scientific elements—including lead optimization, computational methods, chemical intuition, scientific reasoning, hypothesis formation, argument development, and the overarching narrative—were entirely human-generated***. At no point in the current study was AI used to generate, manipulate, or interpret computational data or molecular structures.***

**Importance of Methodological Transparency:**

In contemporary academic writing, it is crucial that the direction of the scientific narrative always remains under the guidance of human understanding. Furthermore, any AI-generated output must be subjected to rigorous scrutiny and fundamentally assumed incorrect until verified through human-derived knowledge. These principles must be applied regardless of the extent of AI usage. ***By transparently disclosing these specific boundaries of AI assistance—no matter how minimal—authors actively reaffirm these principles, demonstrating that the overarching narrative remains entirely in human hands.***

**AI-Assisted Manuscript Copyediting**

*The following AI-assisted steps were utilized during the copyediting process:*

**Phase 1: Structural and Stylistic Modifications:** Following the completion of a human-authored first draft, Gemini 3.1 was used to identify inconsistencies in formality, optimize paragraph structure, and enhance overall clarity. The adoption of any AI-proposed modifications was highly selective, only permitting changes that explicitly preserved the original writing style and scientific voice. ***Permissible modifications were strictly limited to refining specific vocabulary or optimizing sentence arrangements within an existing paragraph, ensuring the core narrative remained unaltered. Any AI-generated suggestions that attempted to alter the scientific meaning, introduce unprompted concepts, or overwrite the original writing style were categorically rejected.***

**Phase 2: Logical Clarity Assessment:** To ensure clear logical progression, specific written arguments were graded using Gemini 3.1. This process served strictly as a supplementary clarity check, operating in tandem with informal peer feedback to guarantee that the human-generated logic was clearly communicated. The model was prompted to identify logical leaps or confusing transitions in the writing. ***As with Phase 1, the model was strictly prohibited from generating novel scientific assumptions, logical progressions, or alternative hypotheses during this review.***

**Example Prompts Used in the Copyediting Process**

**Example 1:** *Structural and Stylistic Refinement (Phase 1)* "Review the following paragraph and suggest 1-2 minor wording tweaks and sentence rearrangements to improve formality and clarity. Show how the revised paragraph would look and bold where each proposed edit was implemented. Explain the reasoning behind each suggested change. [Insert Draft Text]"

**Example 2**: *Logical Clarity Assessment (Phase 2)* “I am arguing that the thermodynamic reliance on narrow, highly localized ‘keystone’ interactions of current paradox-breaking drug candidates creates an evolutionary vulnerability that threatens to render them ineffective over time. Grade the logical progression of the following paragraph: identify any logical leaps or confusing transitions in my reasoning. Do not generate any revisions—simply bold the areas that can be improved. [Insert Draft Argument]”

**Example 3:** *Consistency and Brevity Review* “Identify areas in this methodology section where the phrasing is redundant or lacks precision. Provide specific suggestions to tighten the prose to meet standard biochemical journal formatting without modifying, adding, or removing any of the actual experimental steps. [Insert Draft Text]”
